# Magnetic Protein Aggregates Generated by Supramolecular Assembly of Ferritin Cages - A Modular Strategy for the Immobilization of Enzymes

**DOI:** 10.1101/2024.09.13.612799

**Authors:** Gizem Ölçücü, Bastian Wollenhaupt, Dietrich Kohlheyer, Karl-Erich Jaeger, Ulrich Krauss

**Affiliations:** Institute of Bio- and Geosciences IBG-1: Biotechnology, Wilhelm Johnen Strasse, Forschungszentrum Jülich GmbH, D-52425 Jülich, Germany; Institute of Molecular Enzyme Technology, Heinrich Heine University Düsseldorf, Forschungszentrum Jülich GmbH, Wilhelm Johnen Strasse, D-52425 Jülich, Germany; Department of Biochemistry, University of Bayreuth, 95447 Bayreuth, Germany

**Keywords:** enzyme immobilization, protein-protein interactions, protein aggregates, biocatalysis, magnetic protein aggregates, MPAs, CatMPAs

## Abstract

Efficient and cost-effective immobilization methods are crucial for advancing the utilization of enzymes in industrial biocatalysis. To this end, *in vivo* immobilization methods relying on the completely biological production of immobilizates represent an interesting alternative to conventional carrier-based immobilization methods. In this contribution, we present a novel immobilization strategy utilizing *in vivo* produced, magnetic protein aggregates (MPAs). MPA production is facilitated by the expression of gene fusions consisting of genes encoding for the yellow fluorescent protein variant citrine and variants of the iron storage protein ferritin, including a magnetically enhanced ferritin mutant from *Escherichia coli.* Expression of the gene fusions allows supramolecular assembly of the fusion proteins *in vivo*, which is driven by citrine-dependent dimerization of ferritin cages. Upon cell lysis, the assemblies coalesce in solution to form MPAs. The fusion of the mutant *E. coli* ferritin to citrine yields fluorescent, insoluble protein aggregates that display magnetic properties, verified by their attraction to neodymium magnets. We further demonstrate that these novel, fully *in vivo* produced protein aggregates can be magnetically purified without the need for *ex vivo* iron-loading. Utilizing a bait/prey strategy, MPAs were functionalized by the post-translational attachment of an alcohol dehydrogenase to the MPA particles to enable proof-of-concept for enzyme immobilization, giving rise to catalytically-active magnetic protein aggregates (CatMPAs). The resulting (Cat)MPAs could easily be obtained from crude cell extracts via centrifugation, or purified using magnetic columns, and exhibited superior stability. The strategy presented here therefore represents a highly modular method to produce magnetic enzyme immobilizates which can be obtained with high purity.

## 1. Introduction

Enzyme immobilization is a vital technology to obtain reusable and stable biocatalysts with improved properties for industrial application, while remedying the shortcomings of enzymes at the same time; namely, low tolerance to harsh process conditions, stability issues or inhibition of activity (Sheldon and van Pelt, 2013, Datta et al., 2013). To this end, various conventional enzyme immobilization methods exist (Liu, 2020, Mohamad et al., 2015, Homaei et al., 2013), such as physical entrapment where the enzyme of interest is trapped within a membrane or a polymer matrix (Sheldon and van Pelt, 2013), surface immobilization where the enzymes are physically adsorbed onto or covalently linked to the surface of suitable support materials (Barbosa et al., 2015, Bilal et al., 2019), and cross linking (Sheldon, 2011, Jegan Roy and Emilia Abraham, 2004), based on precipitating the proteins from the solution into aggregates (or crystals), followed by cross-linking with a bifunctional reagent. However, these strategies also suffer from various drawbacks such as lowered specific activities, leaching of the enzyme from the support material, high costs associated with carriers and immobilization onto/into such materials, along with labor intensiveness and lack of generalizability (Sheldon and van Pelt, 2013, Wang et al., 2009, Sheldon, 2007, Wahab et al., 2020). Therefore, in recent years, a multitude of alternative, solely biologically based, *in vivo* enzyme immobilization methods have been developed (Ölçücü et al., 2021, Rehm et al., 2016). These methods, relying on various principles include, amongst others, the display of target proteins on polyhydroxyalkanoate biopolymers generated *in vivo (Wong et al., 2020, Rasiah and Rehm, 2009)*, trapping target proteins within biologically produced protein crystals *(Heater et al., 2018)*, generating liquid and hydrogel-like protein condensates based on liquid-liquid phase separation principles (Dzuricky et al., 2020, Wei et al., 2020), or the production of catalytically-active inclusion bodies (CatIBs) (Garcia-Fruitos et al., 2012, Jäger et al., 2020, Diener et al., 2016, Köszagová, 2020). The latter concept requires the fusion of aggregation-inducing peptides/proteins/protein domains to a target protein, resulting in the pull-down of active, correctly folded target within an inclusion body matrix formed by misfolded fusion protein species. All of the aforementioned methods offer numerous benefits from an application point of view, as they do not require the use of additional carrier materials or expensive and time-intensive purification of the target enzyme, and typically yield the desired enzyme immobilizate in one step, directly during heterologous overexpression of the corresponding gene fusions. Therefore, using self-aggregating/segregating proteins that partially retain their functionality and can easily be isolated after cell lysis is a highly desired property for potential applications in biotechnology, prompting the need for further developments in the field. To address this shortcoming, we therefore aimed at obtaining magnetic enzyme immobilizates by solely biological means, without the need for *in vitro* iron loading.

Ferritins are a family of ubiquitous, iron-sequestering proteins, which are readily exploited for a wide range of biotechnological applications due to their ability to store iron, high chemical and thermal stability, self-assembling properties and biocompatibility (Arosio et al., 2009, He and Marles-Wright, 2015, Truffi et al., 2016). Applications of ferritin include, but are not limited to, serving as a contrast agent for imaging (Wang et al., 2011, Jin et al., 2014), vessel for drug delivery through encapsulation of target molecules (Chiou and Connor, 2018), or in the synthesis of semiconductor nanoparticles (Yamashita et al., 2010). While there are examples of enzyme immobilization where the target enzyme is covalently crosslinked onto (Turan, 2018), or encapsulated within ferritin cages (Bulos et al., 2021, Li et al., 2019), or chemically-loaded magnetoferritin being used for immobilization utilizing the E-coil/K-coil protein-protein interaction (Zhang et al., 2019), the currently available methods either lack general applicability, do not utilize the potential magnetic properties of ferritin, or rely on *in vitro* iron loading to confer ferritin with magnetism. For instance, encapsulation by ferritin requires either the fusion of another highly positively charged protein to the target, such as GFP(+36), to direct the encapsulation of the enzyme inside the negatively charged ferritin cavity, or intense genetic modification of the target enzyme to confer it with a highly positive surface charge (Bulos et al., 2021, Li et al., 2019). Further, the small size of the ferritin core allows only limited cargo recruitment (up to 2 enzymes/ferritin cage) and the method does not yield magnetic immobilizates. Similarly, an approach relying on the decoration of ferritin cages with a target enzyme using E-/K-coils requires multiple chromatographic purification steps to obtain E-coil bearing ferritins and K-coil and His-tag bearing target enzyme, where it is necessary for the target to tolerate modifications at both termini (Zhang et al., 2019). Here, additional immobilization of the ferritin-enzyme complex onto an affinity matrix is also necessary due to the soluble nature of the enzyme decorated ferritin cages, where the immobilizates reportedly displayed magnetic properties upon *in vitro* iron loading, despite their aforementioned solubility. Therefore, to the best of our knowledge, there is currently no entirely biological, modular strategy for the generation of magnetic enzyme immobilizates utilizing ferritin, which allows immobilizate purification via magnets without *ex vivo* iron loading.

Furthermore, as the currently established *in vivo* immobilization methods suffer from the disadvantage of requiring the generation and characterization of numerous constructs consisting of the fusions of the target genes to those of aggregating (Jäger et al., 2020, Ölçücü et al., 2021, Rehm et al., 2016), bio-polymer forming (Grage et al., 2009), or liquid-condensate forming proteins (Schuster et al., 2018), which is a time consuming and laborious process, we envisioned to circumvent these shortcomings by generating a self-assembling and aggregating protein scaffold consisting of ferritin and a fluorescent reporter protein. Our approach therefore has been designed to offer the following benefits: i) easy detection of the aggregation efficiency of the scaffold due to the presence of the fluorescent reporter protein, ii) modularity, due to the presence of a protein interaction motif included in the ferritin-based scaffold, which can be easily decorated with soluble target enzymes containing the complementary motif, and iii) two different possibilities to obtain the immobilizates; quickly by simple centrifugation, or with a higher purity via magnetic purification.

To this end, we utilized a previously described fusion protein based on the heavy chain of human ferritin (HuftnH) and the yellow fluorescent protein variant citrine that self-assembles into supramolecular complexes *in vivo*, showing sustained self-aggregation and sedimentation upon cell lysis (Bellapadrona and Elbaum, 2014, Bellapadrona et al., 2015). Firstly, we generated and assessed the magnetism of the Citrine-HuftnH fusion first described by Bellapadrona *et al*., which did not display substantial magnetic properties in our hands. To implement the envisioned immobilization strategy, we therefore exchanged the HuftnH with various ferritins to generate fully biologically produced, magnetic protein aggregates (MPAs). MPAs containing different ferritins (Figure 1, Panels A and C) were characterized with regard to their aggregation efficiencies and magnetic properties, and the best performing fusion construct was subsequently used to generate magnetic enzyme immobilizates by using the SpyTag/SpyCatcher protein conjugation system (Zakeri et al., 2012, Zakeri and Howarth, 2010) to allow immobilization of an alcohol dehydrogenase model enzyme, resulting in catalytically-active magnetic protein aggregates (CatMPAs, Figure 1, panels D and E) decorated with the target enzyme. Furthermore, two different approaches to isolate the MPAs directly from the crude cell extracts are developed, which offer the benefit of quick and easy purification via centrifugation, or obtaining high purity via magnetic columns. The ferritin-based CatMPAs therefore represent a modular, novel scafold to immobilize enzymes, applicable for both *in vivo* and *ex vivo* enzyme immobilization, and thus can serve as a promising new tool for biotechnological applications in the future.

**Figure 1.**
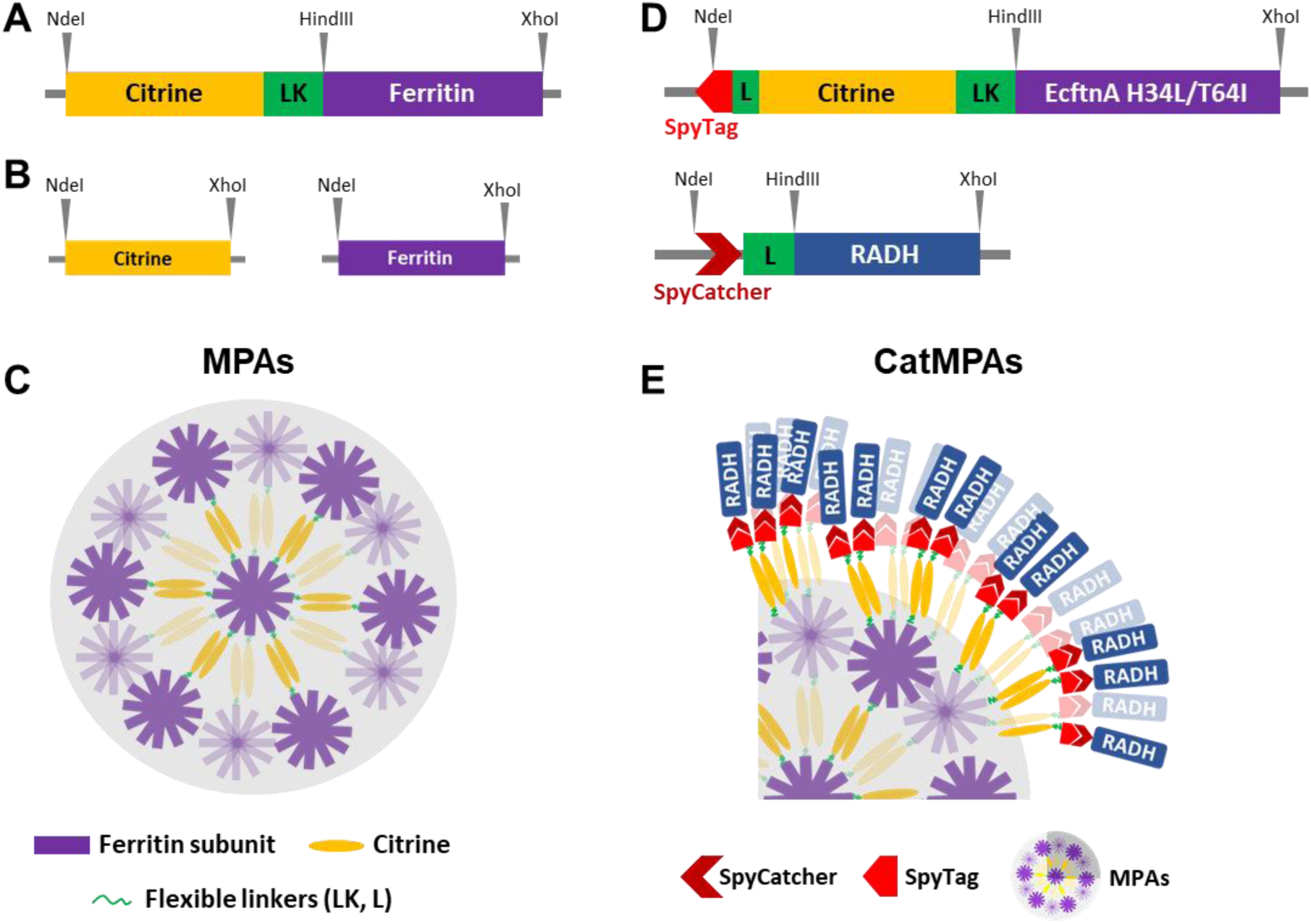
Constructs generated in this study and cartoon diagram depicting the supramolecular assembly of MPAs and CatMPAs. **(A)** Depiction of citrine-ferritin constructs flanked by NdeI and XhoI sites, where a 17-residue linker (LK) with the amino acid sequence GGTGGSGGSGGSGGTGG followed by the HindIII site separates the genes encoding citrine and ferritin. The flexible linker (L) present in CatMPA constructs is similar to the LK linker of MPA, however has the following amino acid sequence: (GGGGS)2. Ferritin refers either to the heavy chain of human ferritin (HuftnH), the nonheme ferritin from *E. coli* (EcftnA-WT) or the double mutant of the *E. coli* ferritin (EcftnA H34L/T64I). **(B)** Depiction of the soluble citrine and ferritin constructs flanked by NdeI and XhoI sites. **(C)** Simplified cartoon drawing of the supramolecular assembly of MPAs. Ferritin subunits self-assemble to form the ferritin cage, and the citrines attached to each ferritin subunit form dimers, giving rise to the depicted supramolecular assembly. For simplicity, only half of the ferritin subunits are shown in the cartoon diagram. **(D)** Depiction of the SpyTag/SpyCatcher bearing constructs used in the CatMPA approach. The gene encoding the SpyTag fragment is present at the 5’ of the gene encoding Citrine-EcftnA H34L/T64I fusion in the bait construct, and the gene encoding the SpyCatcher fragment is present at the 5’ of the gene encoding an alcohol dehydrogenase (RADH) for the prey construct. For both SpyTag and SpyCatcher bearing constructs, a flexible (GGGGS)2 linker (L) separates SpyTag/SpyCatcher from the remaining gene fusion, whereas the flexible linker (LK) with the amino acid sequence GGTGGSGGSGGSGGTGG is present between citrine and ferritin, similar to MPA constructs. **(E)** CatMPAs produced by mixing the bait and prey fusion proteins depicted in panel (D). The MPAs formed by the SpyTag-Citrine-ferritin (bait) construct are decorated by the SpyCatcher-RADH fusion protein due to the interaction of the SpyTag/SpyCatcher pair. For simplicity of the illustration, only a quarter of the MPA particles depicted in panel (C) is shown, and only some of the ferritin subunits are shown to interact with the prey via the citrines, whereas theoretically all 24 ferritin subunits are fused to a citrine bearing the SpyTag which can recruit prey. See methods for additional information and the cloning procedure, and Table S1 for the list of all constructs generated in the study.

## 2. Materials and methods

### 2.1 Generation of gene fusions and expression constructs

For the generation of Citrine-HuftnH (Bellapadrona et al., 2015), Citrine-EcftnA-WT and Citrine-EcftnA H34L/T64I (Liu et al., 2016) constructs, synthetic genes encoding the fusion proteins flanked by 5’-NdeI and 3’-XhoI sites were synthesized (Invitrogen GeneArt Gene Synthesis, ThermoFischer Scientific). All constructs contained a flexible linker (LK) harboring a 3’-HindIII site between the genes encoding citrine and ferritins. Additionally, since the *ecftna* gene naturally encodes an NdeI site (nucleotides 157-162), the thymine at position 159 was exchanged to cytosine during the design of the genes to simplify the cloning process. Therefore, all EcftnA-WT and EcftnA H34L/T64I constructs generated in this study contained this silent mutation. The plasmids harboring the synthetic genes were hydrolyzed with NdeI and XhoI restriction endonucleases, and were ligated into similarly hydrolyzed pET28a (Merck, Darmstadt, Germany) which was used as expression plasmid. A control strain for the production of soluble citrine lacking ferritin was generated via PCR by employing suitable oligonucleotide primers with 5’-NdeI and 3’-XhoI sites (Table S2), using the Citrine-HuftnH construct as template. The resulting PCR product was digested with NdeI and XhoI, and ligated into similarly hydrolyzed pET28a. For the generation of SpyTag002 (Keeble et al., 2017) and SpyCatcher002 (Keeble et al., 2017, Keeble and Howarth, 2019) (optimized variants of SpyTag and SpyCatcher, respectively, referred simply as SpyTag/SpyCatcher in the manuscript) bearing strains, the DNA fragments encoding the SpyCatcher, SpyTag and a (GGGGS)2 linker (L) sequence were synthesized (Invitrogen GeneArt Gene Synthesis, ThermoFischer Scientific). The bait construct SpyTag-Citrine-EcftnA H34L/T64I was generated by hydrolyzing the synthetic SpyTag-Citrine gene fusion flanked by 5’-NdeI and 3’-HindIII sites, and ligating the resulting fragment to similarly digested Citrine-EcftnA H34L/T64I containing pET28a vector, and contained the linker (L) sequence separating the gene encoding SpyTag from the gene fusion encoding citrine-ferritin. The prey construct SpyCatcher-RADH was generated by the amplification of the synthetic SpyCatcher sequence using primers with 5’-NdeI and 3’-HindIII sites (Table S2), followed by hydrolyzing the PCR product by these restriction enzymes, and ligating it to similarly hydrolyzed vector containing the RADH sequence that was generated elsewhere (Ölçücü et al., 2022).The SpyCatcher-RADH construct hence contained a cleavage site for the Factor Xa protease followed by a HindIII site at the 3’ end of the linker (L) separating the genes encoding SpyCatcher and RADH). All constructs were verified by sequencing (Seqlab GmbH, Göttingen, Germany).

### 2.2 Bacterial strains, media and cultivation

*E. coli* DH5α served as the cloning host for the generation of the constructs. For heterologous expression, *E. coli* BL21(DE3) was used. Lysogeny broth (Bertani, 1951) served as the growth medium for the cultivation during cloning and for the precultures for heterologous overexpression of the gene fusions. Autoinduction(AI) medium (Studier, 2005) (12 g/l casein-hydrolysate, 24 g/l yeast extract, 2.2 g/l KH2PO4, 9.4 g/l K2HPO4, 5 g/l glycerol at pH 7.2 supplemented with 0.5 g/l glucose and 2 g/l lactose) was used as the growth medium during protein production. 50 μg/ml kanamycin was added to all growth media for plasmid maintenance. Briefly, LB precultures were used to inoculate the expression cultures with an initial OD600 of 0.05 and were cultivated at 37 °C for 3 hours, shaking at 130 rpm. After 3 hours, the ferritin containing strains were supplemented with iron-citrate complex to a final concentration of 1 mM (0 - 10 mM iron-citrate for BioLEctor experiments with varying iron concentrations, Figure S2), using a sterile filtered stock solution of 100 mM FeSO4·7H2O-500 mM citrate pH 7, and all expression cultures were transferred to 15 °C for 69 hours at 130 rpm. For microscopy, soluble citrine and Citrine-HuftnH/EcftnA-WT/EcftnA H34L/T64I strains were cultivated in a BioLector setup in M9-AI medium (5 g/l (NH4)2SO4, 3 g/l K2HPO4, 6.8 g/l Na2HPO4, 0.5 g/l NaCl, 2 g/l NH4Cl, 0.2 g/l MgSO4·7H2O, 1.5 mg/l CaCl2· 5H2O, 15 mg/l FeSO4, 0.2 g/l Na3C6H5O7·2H2O, 10 mg/l thiamine, 0.75 mg/l AlCl3·6H2O, 0.6 mg/l CoCl2·6H2O, 2.5 mg/l CuSO4·5H2O, 0.5 mg/l H3Bo3, 17.1 mg/l MnSO4·H2O, 3 mg/l Na2MoO4·2H2O, 1.7 mg/l NiCl2·6H2O, 15 mg/l ZnSO4·7H2O, 5 g/l glycerol, 0.5 g/l glucose and 2 g/l lactose) supplemented with 1 mM iron-citrate and were inoculated at a starting OD600 of 0.05 from LB precultures grown overnight. The initial cultivation was performed at 37 °C for 3 hours shaking at 1200 rpm, and expression took place at 15 °C for 69 hours at 1200 rpm, after which the live cells were imaged (see 2.8)

### 2.3 Preparation of cell fractions

*E. coli* BL21(DE3) cells overproducing the target proteins or protein fusions were harvested by centrifugation (6500xg, 30 min, 4 °C). Cells were resuspended 10% (w/v) in lysis buffer (50 mM sodium phosphate buffer, 100 mM NaCl, pH 7.0 for SpyTag/SpyCatcher bearing CatMPA constructs, pH 8.0 for the remaining constructs). For CatMPA constructs, the lysis buffer at pH 7.0 also served as the incubation buffer for the SpyTag-SpyCatcher reaction to take place. Cells were lysed by using an Emulsiflex-C5 high pressure homogenizer (Avestin Europe GmbH, Mannheim, Germany) with internal pressure between 1000-1500 bar, 3 cycles under constant cooling. For CatMPA constructs, the freshly obtained crude cell extracts (CCEs) of bait and prey were mixed in 1:1 (v/v) ratio, vortexed for a few seconds, and then incubated at 25 °C for 30 minutes with shaking at 600 rpm. After 30 minutes, the mixed CCEs were placed on ice. To obtain the soluble and insoluble cell fractions, fresh CCE (or the CCE mixture of bait+prey) was diluted using lysis buffer, and half of the diluted CCE was centrifuged for fractionation (7697x*g*, 2 min, room temperature) as described elsewhere (Ölçücü et al., 2022). The supernatant (S) was transferred to a fresh tube and the unwashed pellet (P1) was resuspended using the same volume of lysis buffer as the removed S fraction. The suspended pellet was centrifuged (7697x*g*, 2 min, room temperature), and the supernatant of the wash (S2) was discarded. The washed pellet was resuspended again in the same volume of lysis buffer as the removed supernatant, resulting in the washed pellet fractions (P). The obtained cell fractions (CCE, S and P) were subsequently kept on ice and were used to determine the fluorescence/RADH activity distributions of the constructs and their mixtures. The MPA pellets were further lyophilized as described elsewhere ((Ölçücü et al., 2022), see also SI methods) to determine yields, and the respective CatMPA and bait-prey pellets were lyophilized similarly for stability analyses.

### 2.4 Imaging over permanent neodymium magnets

To visualize the magnetic properties of citrine-ferritin fusions, 5 ml of crude cell extracts (CCE) of constructs overproducing the citrine-ferritin fusions were mixed with 1 ml of OptiPrep Density Gradient Medium (STEMCELL Technologies Germany GmbH, Köln, Germany) and transferred to mini petri dishes, corresponding to 10% iodixanol (w/v) concentration in the mixture. The CCE-OptiPrep mixture was supplemented with 50 μg/ml kanamycin to prevent contamination during the course of imaging. Four permanent, axially magnetized N45 neodymium ring magnets (with the dimensions of 20 mm (outer diameter), 10 mm (inner diameter), 6 mm height, EarthMag GmbH, Dortmund, Germany) were arranged in a 2x2 grid and were used to assess the magnetic properties of the constructs visually. Black papers cut in a rectangular shape were placed over the neodymium magnets to aid visualization in a similar way as described elsewhere (Liu et al., 2016, Nishida and Silver, 2012), and the mini petri dishes containing the CCEs were placed carefully on top of the papers resting over the neodymium magnets. The samples were imaged every 10 minutes for up to 69 hours, and the patterns emerging in the solution due to the attraction of the citrine-ferritin particles towards the neodymium magnets were captured using a camera (Logitech C930E Full HD-Webcam, Logitech Europe S.A., Lausanne, Switzerland) which was placed directly above the samples. The time lapse video was created using images taking at 10 minute intervals via SkyStudioPro, and edited using DaVinci Resolve 17 (Blackmagic Design Pty Ltd., Melbourne, Australia) to minimize flickering.

### 2.5 Magnetic column purification

To magnetically purify ferritin fusion proteins, commercial MS columns containing ferromagnetic spheres were placed in an OctoMACS separator containing a permanent magnet, held by a MACS multistand (Miltenyi Biotec B.V. & Co. KG, Bergisch Gladbach, Germany). The crude cell extracts of MPAs and CatMPAs were supplemented with 0.05 mg/ml DNase I to prevent clogging of the MS columns prior to application. The lysis buffer (50 mM sodium phosphate buffer, 100 mM NaCl, pH 7.0 for SpyTag/SpyCatcher constructs and pH 8.0 for the remaining constructs) was degassed to get rid of air bubbles that could likewise clog the column. 1 ml of degassed lysis buffer was used to wet the MS column placed in an OctoMACS separator and the eluate was discarded. After the equilibration step, 1 ml of CCE was passed through the MS column and collected, and the eluted CCE sample was reloaded onto the same MS column for a total of three times. The sample that eluted after the third run was collected and labelled as the NM (nonmagnetic) fraction. The column was then washed two times using 1 ml degassed lysis buffer and the eluates were collected separately as wash fractions W1 and W2. To obtain the MG (magnetic) fraction, the MS column was removed from the OctoMACS separator, loaded with 1 ml degassed lysis buffer, and the magnetic particles suspended in the column were quickly flushed out using the plunger provided in the kit and collected in a separate tube. All fractions were kept on ice until further analysis.

### 2.6 Fluorescence spectrophotometry

Fluorescence emission of cell fractions of the citrine-containing fusions were measured in quadruples using black Nunc 96-Well MicroWell polypropylene plates (ThermoFisher Nunc, Waltham, USA) and a TECAN infinite M1000 PRO fluorescence MTP reader (TECAN, Männedorf, Switzerland). 100 µl of CCE, S, P cell fractions or NM, W1, W2, MG fractions in appropriate dilutions were applied in quadruples onto the microtiter plates and the fluorescence emission of the samples were quantified (λex = 513 nm, λem = 529 nm, z-position 18.909 µm, enhancement 120, flash number 25, flash frequency 400 Hz, bandwidth 5 nm). Samples were shaken (654 rpm, 2 mm amplitude) for 5-10 seconds immediately before the fluorescence measurements to ensure that the particles are suspended. All measurements were performed using at least three biological replicates.

### 2.7 RADH activity measurements

The cell fractions of prey constructs containing RADH, along with the respective fractions of CatMPA CCE mixtures were tested for the distribution of the RADH activity using a discontinuous photometric assay where the consumption of the NADPH was detected as described earlier (Jäger et al., 2019, Ölçücü et al., 2022). Briefly, RADH containing cell/magnetic purification fractions and a reaction mixture of 1400 µl containing 0.5 mM NADPH and 125 mM cyclohexanone in TEA-buffer (50 mM Triethanolamine, 0.8 mM CaCl2, pH 7.5) were incubated separately at 30 °C for 5 minutes. The reaction was initiated by transferring 350 µl of the RADH containing sample onto the 1400 µl reaction mixture, immediately vortexed, and a sample of 250 µl was taken which was transferred onto 500 µl of methanol to stop the reaction. The remaining reaction mixture was quickly placed in a shaking incubator at 1000 rpm and 30 °C. The rest of the reaction mixture was then sampled every minute for a total of six times in the same manner, where the sampled reaction was stopped in methanol. After the last sampling step, the vials were centrifuged (7697*g*, 5 minutes, room temperature) and transferred to disposable cuvettes to measure the absorption spectra (280 - 500 nm) using a Cary 60 UV-Vis Spectrophotometer (Agilent, Santa Clara, USA). All measurements were performed using at least three biological replicates. Stability analyses were performed similar to described elsewhere (Ölçücü et al., 2022). In brief, 14 mg of CatMPA or the SpyCatcher-RADH prey lyophilizates were suspended in 7 ml of lysis buffer (50 mM sodium phosphate buffer, 100 mM NaCl, pH 8.0) and vortexed until no visible clumps remained in the solution. The RADH activity of the lyophilizate suspensions were measured in triplicates as described above for the cell fractions, where the reaction mixture was sampled once every 2 minutes instead of 1 minute, for a total of 10 minute assay, in order to have increased sensitivity. After the assay, the remaining lyophilizate suspensions were placed at a 25 °C incubator for 5 days, where the RADH activity assay was repeated in triplicates every 24 hours.

### 2.8 Microscopic analyses

Live *E. coli* BL21(DE3) cells producing the MPA constructs and soluble citrine were cultivated in M9-AI medium supplemented with 1 mM iron-citrate complex as described above. At the end of expression (69 hours), cultures were diluted appropriately in lysis buffer (50 mM sodium phosphate buffer, 100 mM NaCl, pH 8.0) and 1 µl of the cell suspension was transferred to glass slides and covered with a coverslip. The samples were then analyzed with Nikon Eclipse Ti microscope (Nikon GmbH, Düsseldorf, Germany) with a YFP filter (λex = 500 nm, λem = 542.5 nm) and Nikon DS-Qi2 camera (Nikon GmbH, Düsseldorf, Germany). Fluorescence and camera exposure times were 200 ms for ph3 and 100 ms for the YFP filter used to detect citrine fluorescence. For CatMPA constructs, approximately 1 ml was sampled at the end of expression, centrifuged (7697g, room temperature, 1 minute), resuspended and diluted suitably using lysis buffer. The cell suspension was then transferred to polydimethylsiloxane microfluidic chips with inner chamber dimensions of 60 µm x 100 µm x 1µm, and imaged using Nikon Nikon Eclipse Ti microscope (Nikon GmbH, Düsseldorf, Germany) with a YFP filterblock (λex 495 nm, λem 520 nm) and Andor Zyla VSC-01418 camera (Oxford Instruments plc, Oxon, UK) with exposure times of 100 ms for ph3 and YFP filters.

### 2.9 Determination of protein concentration and SDS-PAGE analyses

Protein concentration of the supernatant samples (S) were determined with the Bradford assay(Bradford, 1976) and bovine serum albumin standards with concentrations between 0.01 - 0.1 mg/ml. Either home-made SDS-gels (5-12%) or NuPAGE 4-12% Bis-Tris protein gels in MES SDS running buffer (50 mM MES, 50 mM TRIS, 0.1% SDS, 1 mM EDTA, pH 7.3) were used for SDS-PAGE analyses. The volume required to have 10 µg of protein based on the Bradford assay for the S fraction was set as the loading volume for the remaining cell and magnetic purification fractions except for MG. For the MG fraction, the sample was applied onto polyethersulfone membrane centrifugal filters with 3 kDa cutoff (VWR International GmbH, Darmstadt, Germany) to concentrate this fraction. The concentrated MG sample was loaded onto the SDS gel to contain 20 µg of protein in order to increase sensitivity to impuritites. Cell fractions were boiled at 100 °C for 3 minutes before loading onto the SDS gels, and each gel contained 3 µl PageRuler Prestained Protein Ladder (ThermoFisher Nunc, Waltham, USA).

## 3. Results and Discussion

### 3.1 Diversification of a ferritin-based self-assembly system

To obtain biologically produced, magnetic immobilizates, we initially reconstructed a fusion protein consisting of the fluorescence reporter citrine and the heavy chain of human ferritin (Citrine-HuftnH) as first described by Bellapadrona and co-workers (Bellapadrona et al., 2015, Bellapadrona and Elbaum, 2014). Citrine-HuftnH had been shown to yield self-assembling supramolecular complexes, producing fluorescent particles in *E. coli*, which further aggregated and sedimented in solution upon release from the cells. Self-assembly and aggregation was postulated to be due to dimerization of citrine attached to the ferritin subunits that themselves assemble to intact ferritin cages, with the citrines mediating the formation of the supramolecular complexes (Bellapadrona and Elbaum, 2014, Bellapadrona et al., 2015) (Figure 1, C). To extend on this strategy, we used a nonheme *E. coli* ferritin (EcftnA-WT), and a magnetically enhanced EcftnA H34L/T64I (Liu et al., 2016) in addition to the HuftnH, and fused the genes encoding the human and *E. coli* ferritins to the 3’ end of the gene encoding citrine (Figure 1, A). For initial assessment of the self-aggregation properties of all constructs, Citrine-HuftnH, Citrine-EcftnA-WT and Citrine-EcftnA H34L/T64I fusions were overproduced in *E. coli* BL21(DE3) and the cells were lysed to yield the crude cell extract (CCE) fractions. All CCEs visually showed self-aggregation and sedimentation when left undisturbed (Figure S1), confirming that the exchange of human ferritin with *E. coli* ferritins did not interfere with the aggregation tendency of the fusion proteins.

The presence of intracellular supramolecular aggregates was further confirmed via microscopic analyses conducted on live *E. coli* BL21 (DE3) cells overproducing the citrine-ferritin fusions, with a construct producing soluble citrine included as a negative control (Figure 2). All citrine-ferritin fusions exhibited localized signals for citrine fluorescence at only one of the cell poles, whereas citrine control construct displayed uniform fluorescence as expected. This observation is also in line with the relative fluorescence data (Figure 3), and the literature on the Citrine-HuftnH construct (Bellapadrona et al., 2015), where Citrine-HuftnH exhibited localized fluorescence signals. It should be noted that the aggregates produced by citrine-ferritins are visually different when compared to conventional (catalytically-active) inclusion bodies (CatIBs) (Jäger et al., 2019, Jäger et al., 2018, Garcia-Fruitos et al., 2005, Ölçücü et al., 2022), as the citrine-ferritin aggregates appear smaller in size and are predominantly present at just one cell pole, as opposed to inclusion bodies which are in general present at both poles. However, the number of generations after induction can have a significant impact on the number and size of the inclusion bodies (IBs) found in bacteria. For instance, spherical IBs are more commonly observed in ‘older’ cell cultures, and once IB size is sufficiently large, they tend to fuse into one larger IB, which is thought to occur via the agglomeration of the rod-shaped, early-stage IBs (Kopp et al., 2023). While the time between induction and harvest is sufficient for the formation of a single, spherical IB in the MPA producing cells in our experiments, the smaller size of the fluorescent signal originating from the single pole in comparison to CatIBs which were imaged under the same conditions (Jäger et al., 2019, Jäger et al., 2018, Garcia-Fruitos et al., 2005, Ölçücü et al., 2022), as well as the lack of refractile particles under phase contrast microscopy (data not shown) which is a typical feature of IBs/CatIBs, indicate that MPAs are likely not IB-based materials.

**Figure 2.**
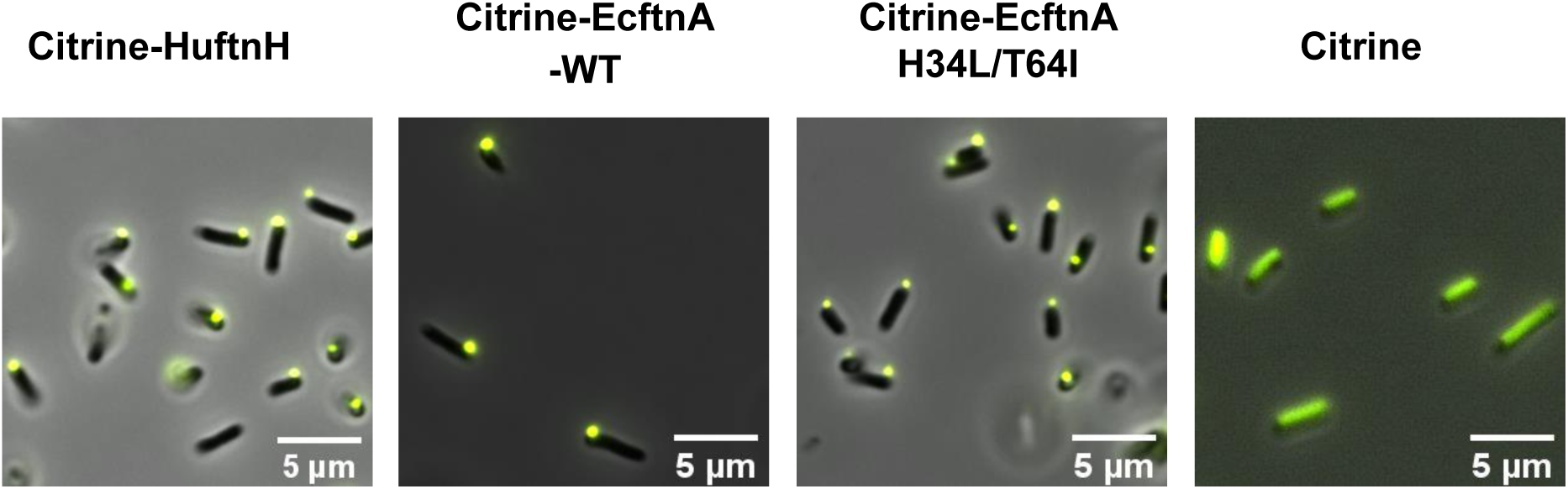
Fluorescence microscopy pictures of live *E. coli* BL21(DE3) cells overproducing citrine-ferritin fusions and soluble citrine. See methods section for details and cultivation conditions.

**Figure 3.**
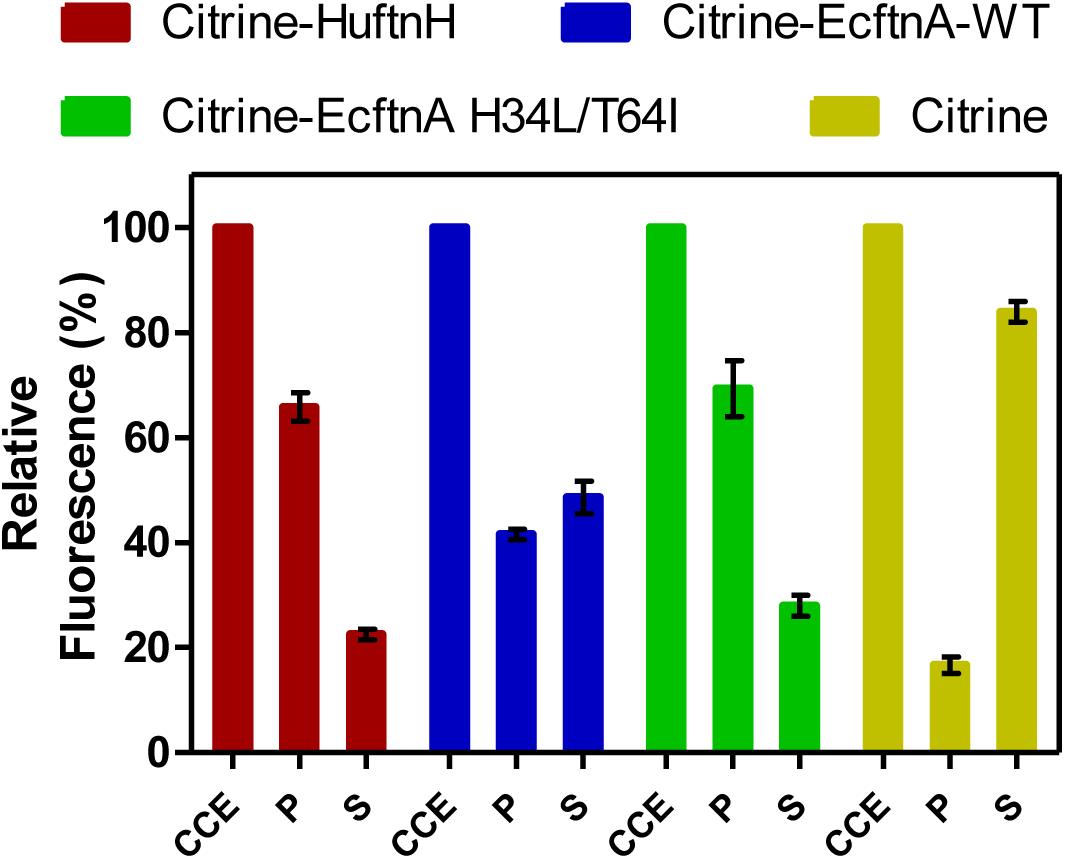
Relative fluorescence of cell fractions from citrine-ferritin fusions and soluble citrine control. Citrine fluorescence of the crude cell extract (CCE) fraction was set to 100% for each construct, and the fluorescence signal detected in washed pellet (P) and supernatant (S) fractions are shown relative to the fluorescence of their corresponding CCE fractions. The error bars represent standard error of the mean derived from at least three biological replicates with four technical replicates each.

To quantify the aggregation efficiencies of all constructs, CCEs were fractionated (see Preparation of cell fractions) by centrifugation to yield the soluble supernatant (S) and the insoluble pellet fractions for all constructs. The pellets were then washed and centrifuged a second time to yield the washed pellet (P) fractions, which allowed the quantification of citrine fluorescence distributions for all constructs (Figure 3). Citrine fluorescence detected in the P fraction was then compared to the fluorescence of the CCE fraction (set to 100%) to assess the aggregation efficiencies (%) for all constructs. A construct overproducing soluble citrine was also included in the analysis as control.

As evident from Figure 3, all pellets obtained from the citrine-ferritin fusions were fluorescent and the Citrine-EcftnA H34L/T64 construct displayed the highest aggregation efficiency among the generated constructs, where 69% of the total fluorescence signal of the CCE originated from the insoluble, washed pellet fraction for this construct. Citrine-HuftnH and Citrine-EcftnA-WT constructs displayed very high aggregation efficiencies as well (66% and 42%, respectively). In contrast, the citrine construct lacking ferritins had only 17% of the citrine fluorescence in the pellet, indicating that in addition to dimerization of citrines, fusion of ferritin to the citrine is crucial for aggregation, which is in line with earlier studies conducted with the HuftnH fusion construct (Bellapadrona et al., 2015, Bellapadrona and Elbaum, 2014). In addition, yields of the constructs were determined, along with their protein contents (Table S4), which indicates that the ferritin-based protein aggregates can be produced at comparable yields (up to 4.7 g lyophilizate / 100 g wet cells, and 77% protein content depending on construct, see Table S4 and SI for the method) with our production and handling techniques as compared to CatIBs (Ölçücü et al., 2022)

In conclusion, we could demonstrate that the HuftnH can be successfully exchanged with *E. coli* ferritins to obtain fluorescent aggregates, and, as evidenced by the case of the EcftnA H34L/T64I mutant, the resulting fusion proteins can exhibit superior aggregation efficiencies.

### 3.2 Magnetic properties of MPAs generated by Citrine-EcftnA H34L/T64I and magnetic purification

After the initial characterization of citrine-ferritin fusions via live cell microscopy and fluorescence spectroscopy of the cell fractions, we investigated the magnetism of citrine-ferritin fusions. To provide an easy, visual indication of the magnetic properties of Citrine-HuftnH, Citrine-EcftnA-WT and Citrine-EcftnA H34L/T64I fusions, the fluorescent citrine-ferritin particles were tested for their response towards permanent magnets, in a similar way that was described elsewhere to test whole cell magnetism (Nishida and Silver, 2012, Liu et al., 2016). To this end, cells overproducing the citrine-ferritin fusions, which were cultivated in autoinduction medium supplemented with 1 mM iron-citrate complex, were lysed (see methods for details and Figure S2 for BioLector experiment with varying iron concentrations). The crude cell extracts (CCEs) of the citrine-ferritin fusions were then transferred to mini petri dishes containing 17% (v/v) OptiPrep density gradient medium. CCE-OptiPrep suspensions were immediately placed over permanent neodymium ring magnets (arrangement shown in Figure 4, leftmost panel) and were imaged up to 69 hours using a camera placed above the samples.

**Figure 4.**
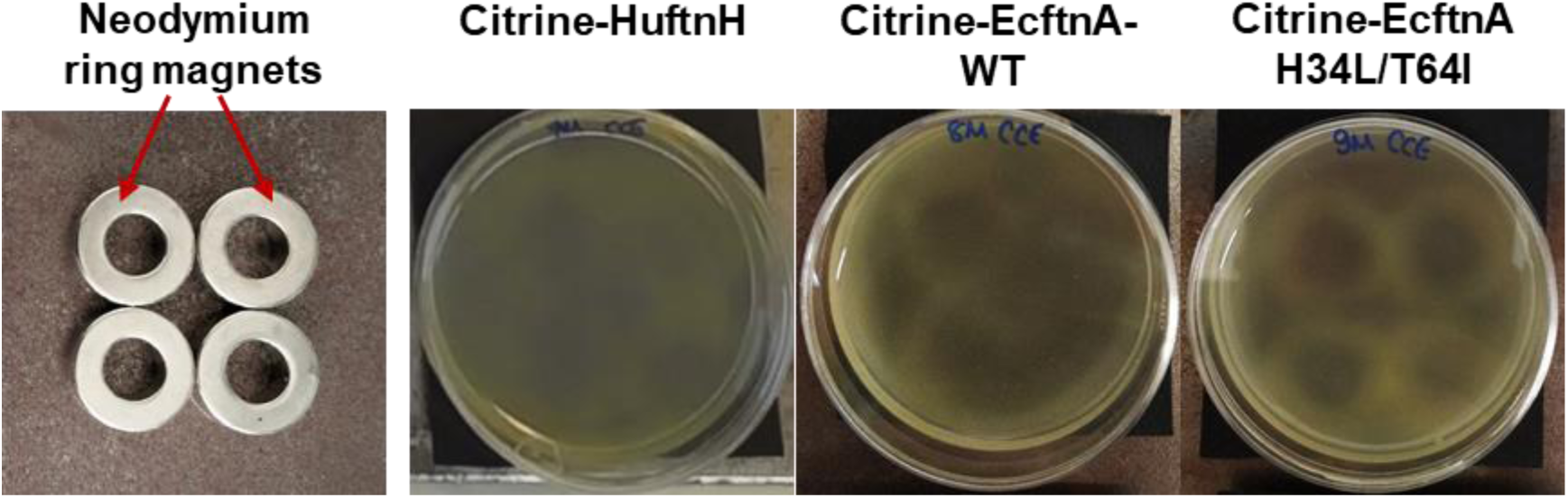
Assessment of the magnetic properties displayed by the crude cell extracts (CCEs) of different citrine-ferritin fusions using permanent neodymium ring magnets. The magnets were arranged in a 2x2 grid as shown in the leftmost panel, and covered with a black paper to aid visualization, onto which the mini petri dishes containing the CCE-OptiPrep density gradient medium mixtures (17% OptiPrep) were placed. Upon placement over the permanent magnets, the CCEs were left undisturbed for up to 69 hours to follow the pattern formation. The contrast of all images shown is increased by 20%.

The attraction of citrine-ferritin particles in the CCE towards the neodymium magnets underneath the suspensions gave rise to patterns of varying intensity for the tested constructs (Figure 4). Faint, albeit noticeable patterns started forming as early as six hours for the CCE of Citrine-EcftnA H34L/T64I construct, and after 12 hours, faint patterns were visible for all three citrine-ferritin constructs (SI video, see Figure S4 for images of citrine-ferritin CCEs visualized every 6 hours). The imaging continued for a total of 69 hours to ensure capturing of the entire pattern progression, which became noticeably sharper for the Citrine-EcftnA H34L/T64I construct as time progressed. As a negative control, cells overproducing HuftnH, EcftnA-WT and EcftnA H34L/T64I without the citrines were cultivated and lysed under identical conditions, and their CCEs were placed over permanent magnets as well, which showed no distinct pattern formation (Figure S3), indicating that the soluble ferritins that lack citrine do not form substantial aggregates.

In conclusion, imaging CCEs of the citrine-ferritin fusions over permanent magnets provided first insights into the magnetic properties of the corresponding MPAs, where the Citrine-EcftnA H34L/T64I construct surpassed Citrine-HuftnH and Citrine-EcftnA-WT constructs in this regard. In addition to showing superior magnetic properties, the Citrine-EcftnA H34L/T64I construct also showed the highest aggregation efficiency as judged by the fluorescence distribution data (Figure 3). Therefore, all further work was conducted using this construct. The magnetic properties of the Citrine-EcftnA H34L/T64I MPAs were further exploited to purify the fusion protein using MS magnetic columns and OctoMACS separator system (Miltenyi Biotec). In brief, the CCE of Citrine-EcftnA H34L/T64I was passed through the same MS column for a total of three times and the eluate was collected (nonmagnetic fraction, NM). The column was then washed twice using lysis buffer (50 mM sodium phosphate buffer, 100 mM NaCl, pH 7.0) and the wash fractions (W1 and W2) were collected. Finally, the magnetic (MG) fraction was eluted by separating the column from the OctoMACS permanent magnet, applying lysis buffer onto the column and quickly flushing the MG fraction using a small plunger. The magnetic column purification fractions were then loaded onto an SDS-PAGE along with the cell fractions obtained via centrifugation, for the assessment of purity of the Citrine-EcftnA H34L/T64I protein.

SDS-PAGE analysis revealed that the Citrine-EcftnA H34L/T64I fusion protein can be purified using magnetic columns, evident by the clear band present in the MG fraction (Figure 5). Furthermore, the washed pellet fraction (P) of the centrifugation approach containing MPAs contained other proteins as well (i.e. possibly chaperons and membrane proteins commonly encountered in CatIB approach (Kloss et al., 2018) for such insoluble fractions), whereas the magnetically purified MPAs were of high purity. Subsequently, the wash fractions of the magnetic purification samples (W1 and W2) were clear, indicating that the columns retain the Citrine-EcftnA H34L/T64I fusion protein rather well, therefore, using magnetic columns appears as a suitable method for purifying MPAs. Furthermore, the citrine-specific fluorescence of the fractions obtained from the magnetic purification approach were determined, and fluorescence detected in each fraction was compared to the total fluorescence of the CCE (set to 100%). Unfortunately, the majority of the citrine fluorescence (approximately 80% of the total CCE fluorescence) originated from the nonmagnetic (NM) fraction, followed by 19% for the magnetic (MG) fraction.

**Figure 5.**
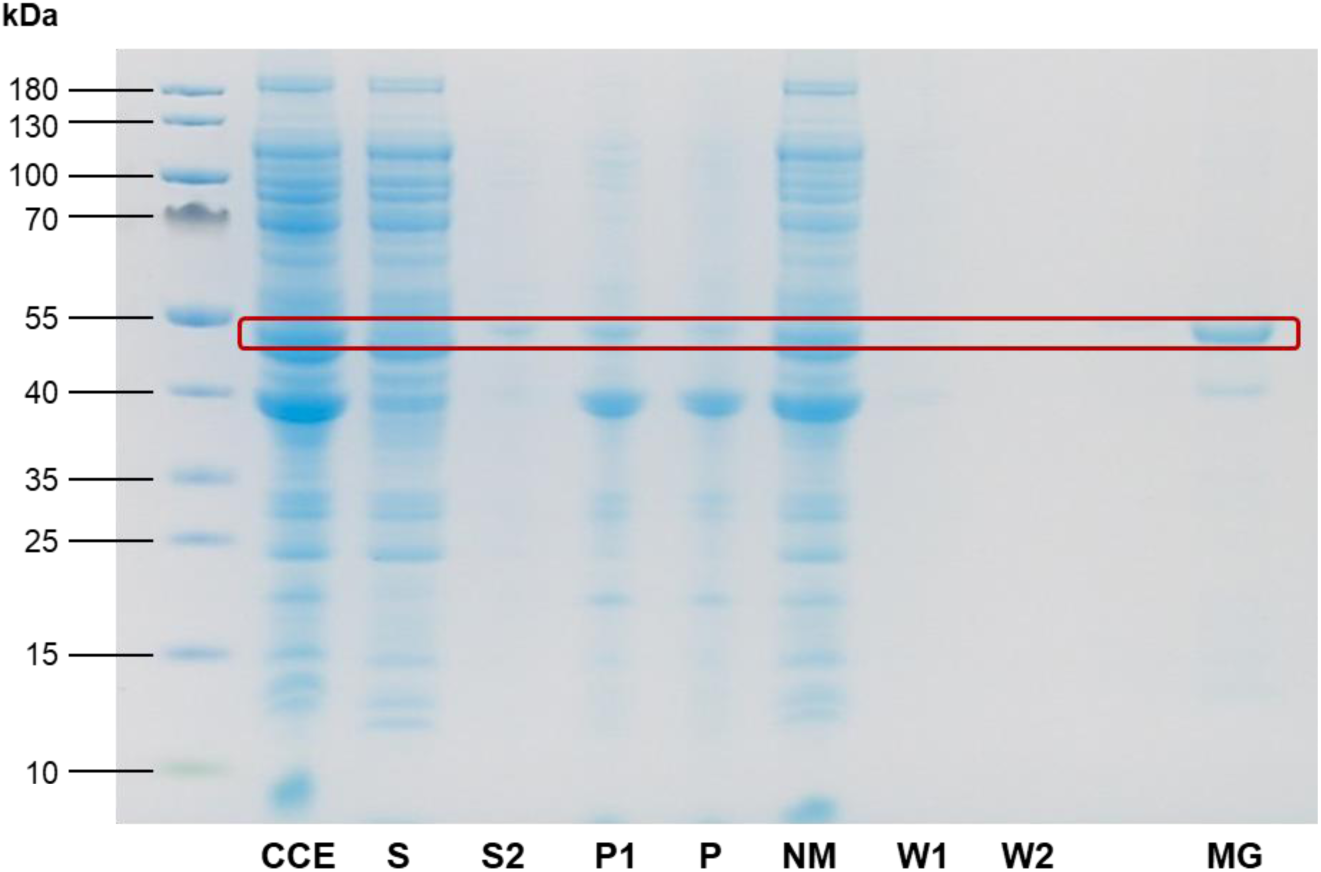
SDS-PAGE analysis of Citrine-EcftnA H34L/T64I cell fractions. The Citrine-EcftnA H34L/T64I fusion protein (47.8 kDa) is marked with a red rectangle for all fractions. CCE: crude cell extract, S(1): supernatant, S2: supernatant of wash step, P1: unwashed pellet, P(2): washed pellet, NM: nonmagnetic fraction, W1: first wash, W2: second wash, MG: magnetic fraction. Protein content of the S fraction was determined using Bradford assay, and the volume required to load 10 µg protein for S fraction was used as the sample volume for all remaining fractions except for MG fraction. The concentration of the MG fraction was determined separately, and the fraction was concentrated prior to loading in order to contain 20 µg protein for this fraction to assess purity of the fraction more critically (See Methods for details, and Figure S5 for SDS-PAGE analyses of cell fractions from all soluble control, MPA, and CatMPA constructs).

The wash fractions W1 and W2 displayed almost no fluorescence (4% and 0.3% when compared to CCE, respectively). As the majority of the fluorescence detected for the Citrine-EcftnA H34L/T64I MPA construct originated from the insoluble fraction (Figure 3), this result indicates that not all of the citrine-ferritin aggregates could be purified by the magnetic purification approach. This could be due to several factors: (i) a significant fraction of citrine-ferritin aggregates exhibits weaker magnetism and are not retained by the column (i.e. due to unequal loading of individual ferritin cages); (ii) the majority of the citrine-ferritin aggregates displays magnetism and are therefore purified, but do not exhibit strong fluorescence. To compare the two approaches quantitatively, we calculated the purification success (%) by comparing the fluorescence of the MG fraction to the fluorescence of the washed pellet (P) fraction obtained via centrifugation (set to 100%). This quantification assumed that all citrine-ferritin aggregates that are obtained via centrifugation could in theory be purified using the columns and would display fluorescence, yielding up to 42% purification efficiency for the magnetic column purification method. Moreover, as the magnetic purification method excludes impurities (Figure 5), it can potentially make up for this loss in cases where high purity is preferable over high quantity.

### 3.3 Extension of the strategy to generate CatMPAs

Next, the magnetic immobilization strategy was further extended as a proof-of-concept to immobilize an alcohol dehydrogenase from *Ralstonia sp.* (RADH) via the CatMPA strategy (Figure 1, E). To this end, we used the SpyTag/SpyCatcher technology (Zakeri et al., 2012), which is based on the engineered CnaB2 domain from a *Streptococcus pyogenes* adhesin, where the SpyTag peptide and SpyCatcher protein arising from the split CnaB2 domain can form a spontaneous, irreversible amide bond that can be used to link two proteins together. We implemented the SpyTag/SpyCatcher system to link the insoluble, Citrine-EcftnA H34L/T64I protein fusion (bait) to soluble RADH (prey), to be able to pull RADH into the insoluble fraction. To this end, the genes encoding SpyTag and SpyCatcher were fused to the 5’ of the genes encoding the bait and the prey, respectively (Figure 1, D). To check if the presence of the SpyTag infers with the generation of fluorescent aggregates for the bait construct, and to confirm that the presence of the SpyCatcher does not result in the formation of significant amounts of RADH inclusion bodies that would shift the RADH to insoluble fraction for the prey, the live cells overproducing both constructs were evaluated by phase contrast and fluorescence microscopy (Figure 6, panels A and C). The microscopic analyses confirmed that the presence of the SpyTag did not interfere with the insoluble fluorescent particle formation for the bait, and SpyCatcher-RADH (prey) showed no particle formation as anticipated.

**Figure 6.**
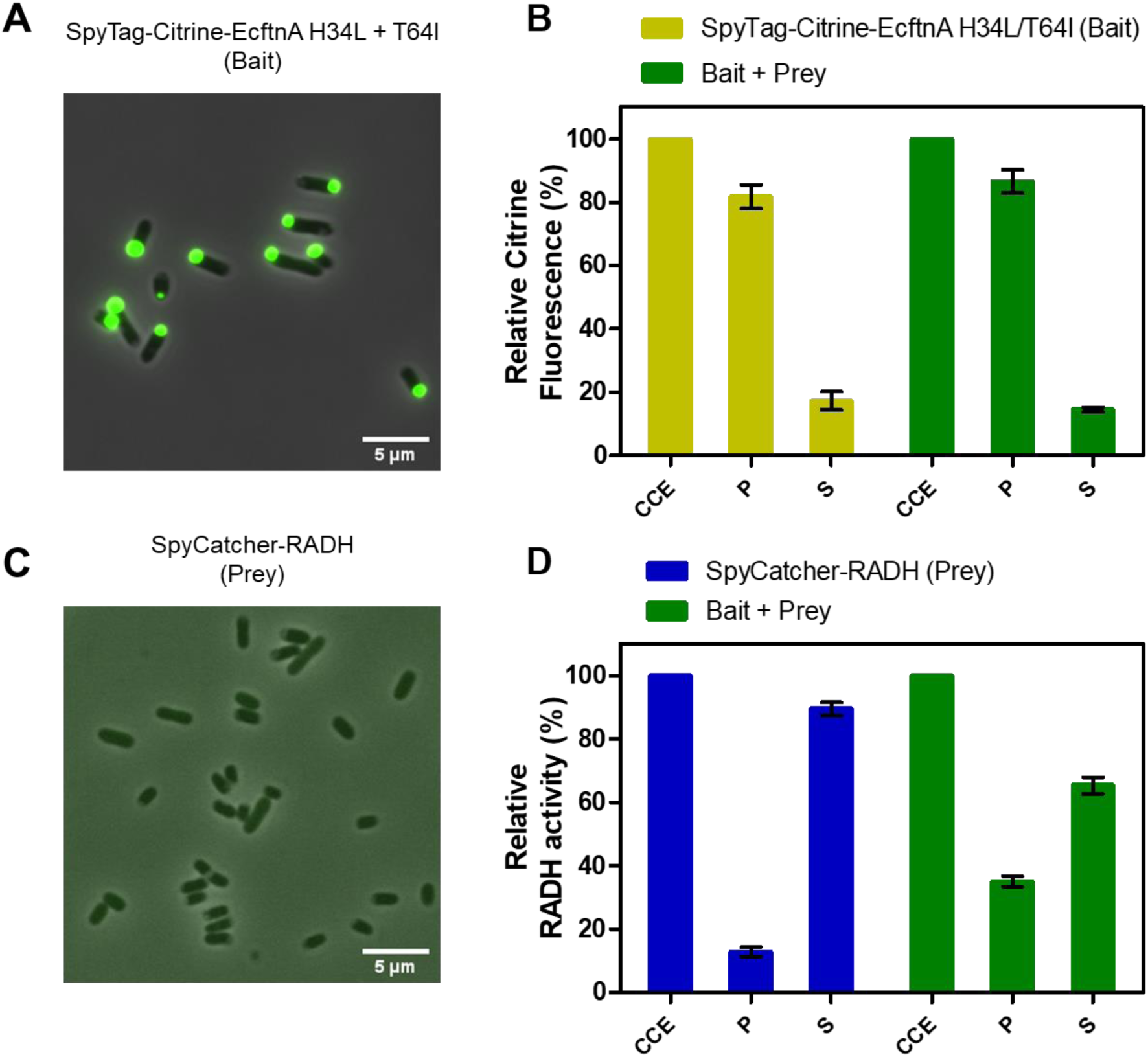
Microscopic analyses and relative fluorescence/activity data for bait and prey constructs. Fluorescence microscopy and phase contrast pictures of live *E. coli* BL21(DE3) cells overproducing SpyTag-Citrine-EcftnA H34L/T64I (bait) is shown in (**A**), and SpyCatcher-RADH (prey) in (**C**). Both panels show composite images obtained by the fluorescence filter and phase contrast. Relative citrine fluorescence (**B**) and relative RADH activity (**D**) of cell fractions of SpyTag-Citrine-EcftnA H34L/T64I (bait, depicted in yellow), SpyCatcher-RADH (prey, depicted in blue) along with the cell fractions of 1:1 (v/v) mixture of the two constructs (bait + prey, depicted in green). CCE: crude cell extract. P: washed pellet. S: supernatant. Error bars correspond to standard error of the mean obtained from at least three biological replicates.

To link the bait and prey constructs, the strains overproducing SpyTag-Citrine-EcftnA H34L/T64I and SpyCatcher-RADH were cultivated separately, the cells were lysed and their CCEs were mixed in 1:1 (v/v) ratio. The CCE mixture was then incubated at 25 °C for 30 minutes to allow the SpyTag/SpyCatcher interaction to take place, after which the mixed CCE was fractionated into soluble and insoluble fractions, and the fluorescence and RADH enzyme activity of the appropriate fractions were determined (for details, see Preparation of cell fractions). The unmixed CCEs of bait and prey constructs were also fractionated to obtain the soluble and insoluble cell fractions, which were tested for fluorescence for the bait construct and RADH activity for the prey (Figure 6). For the bait construct (Figure 6, panel B, yellow bars), citrine fluorescence was detected predominantly in the insoluble fraction (82%), similarly high when compared to the Citrine-EcftnA H34L/T64I construct lacking the SpyTag (Figure 3). For the prey construct, only 13% of the RADH activity could be found in the insoluble fraction (Figure 6, panel D, blue bars). Upon mixing the CCEs of bait and prey constructs, the RADH activity of the insoluble fraction could be increased to 35% of the total RADH activity of the mixture (Figure 6, panel D, green bars), corresponding to almost 3-fold activity increase in this fraction.

Therefore, the RADH activity could be successfully shifted into the insoluble fraction via the SpyTag/SpyCatcher interaction using the CatMPAs. In order to exclude the role of hydrophobic interactions rather than the specific SpyTag/SpyCatcher-based protein-protein interaction, which would also result in an increase of RADH activity for the bait+prey pellet, control experiments were conducted with bait-prey pairs either lacking SpyTag or SpyCatcher. To this end, the CCEs of 2 negative control pairs; i) Citrine-EcFtnA H34L/T64I (bait lacking SpyTag) + SpyCatcher-RADH (prey), and ii) SpyTag-Citrine-EcFtnA H34L/T64I (bait) + soluble RADH (prey lacking SpyCatcher) were mixed and incubated under the same conditions as described for CatMPA generation. The mixed CCEs were subsequently fractionated following incubation as described above. The RADH activity of the pellets derived from these mixtures did not increase significantly when compared to the pellet of the RADH-containing constructs for each case (from 1.66% to 0.89% for the first negative control pair, and from 0.92% to 1.43% for the second pair respectively, see Table S6), which proves the specificity of the protein-protein interactions caused by interaction of the SpyTag/SpyCatcher pair for CatMPA formation.

Furthermore, altering bait:prey ratios and incubation times did not result in significantly higher activity in the insoluble fraction, similar to alternative bait and prey constructs tested which harbored the tags at different termini (Tables S5 and S6). A different approach relying on “magnetization” of GFIL8-RADH CatIBs (Ölçücü et al., 2022) by soluble ferritin cages was likewise tested with different SpyTag/SpyCatcher constructs (Table S6), however, SpyTag-Citrine-EcftnA H34L/T64I and SpyCatcher-RADH combination presented here yielded the best results in terms relative RADH activity for the pellet derived from the bait+prey mixture (Table S6). For instance, the combination of SpyCatcher-EcftnA H34L/T64I and GFIL8-RADH-SpyTag pair, which relied on the magnetization of insoluble GFIL8-RADH CatIBs with soluble ferritin, yielded the highest purification yield (over 97%, Table S6), where these magnetized CatIBs could be purified via magnetic columns, similar to MPAs. However, the here presented SpyTag-Citrine-EcftnA H34L/T64I + SpyCatcher-RADH CatMPA pair showed higher RADH activity and aggregation efficiency. The CatMPAs of SpyTag-Citrine-EcftnA H34L/T64I + SpyCatcher-RADH CatMPA pair could also be magnetically purified, albeit with a lowered purification efficiency (9.4 - 18.4%, as calculated from the relative citrine fluorescence and RADH activity of the magnetic fraction compared to those of the washed pellet, respectively, Table S6) when compared to best performing magnetized CatIBs. The difference in CatMPA purification efficiencies calculated via assays conducted on citrine and RADH indicate different amounts of active protein being present in magnetic and washed pellet fractions, where citrine-ferritins with a higher purification efficiency conversely showing a low citrine fluorescence for the bait fusion. Nevertheless, utilizing a bait-prey approach appears to be a feasible way to generate CatMPAs, and testing different bait-prey constructs and combinations proved beneficial for the optimal implementation of the presented strategy.

Finally, the stability of the CatMPAs was demonstrated by incubating lyophilized CatMPAs, as well as SpyCatcher-RADH, suspended in lysis buffer (50 mM sodium phosphate buffer, 100 mM NaCl, pH 8.0, see methods) at room temperature for 5 days, in the same way as were performed for RADH CatIBs (Ölçücü et al., 2022). Remarkably, the CatMPAs did not lose any activity over a 5 day period, where the RADH activity at the end of 5 days corresponded to 101.3% of the initial RADH activity detected in day 1 (Figure 7). Within the same time period, the activity of the prey dropped to 38.7% of its initial activity, which is comparable to that of soluble RADH (only 31% after 5 days, under the same conditions (Ölçücü et al., 2022)). This demonstrates the remarkable stability of CatMPAs, suggesting that CatMPA-based immobilizates represent a promising new enzyme immobilizate for application in biocatalysis and synthetic chemistry.

**Figure 7.**
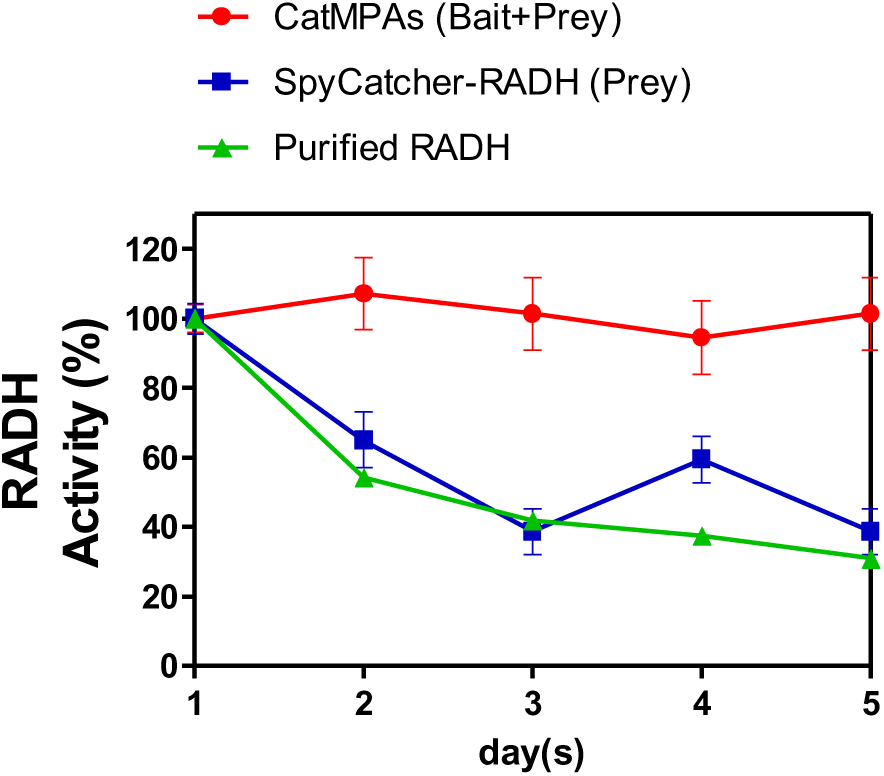
Stability of CatMPAs generated by the SpyTag-Citrine-ferritin (bait) and SpyCatcher-RADH (prey) constructs, compared to prey construct, and soluble, purified RADH over 5 days. Stability of soluble RADH is taken from (Ölçücü et al., 2022). The error bars depict the standard error of the mean from at least three replicates.

## 4. Conclusions

In this study, we successfully generated magnetic protein aggregates (MPAs) by the overproduction of citrine-ferritin fusions, and extended the strategy from human ferritin (Bellapadrona et al., 2015) to wild-type and magnetically enhanced E. coli ferritin (Liu et al., 2016) variants to obtain constructs with superior aggregation efficiencies. Furthermore, we were able to demonstrate the magnetic properties displayed by the MPAs for the first time, and further exploited this property to purify and obtain protein immobilizates of high purity directly from crude cell extracts. To the best of our knowledge, this is the first report describing the generation of fully *in vivo* produced protein aggregates which could be magnetically purified without *ex vivo* iron loading. In proof-of-concept experiments, we generated enzyme-linked magnetic aggregates utilizing the SpyTag/SpyCatcher (Keeble et al., 2017, Keeble and Howarth, 2019) technology to link citrine-ferritin MPAs to an alcohol dehydrogenase (RADH) and produced catalytically-active magnetic protein aggregates (CatMPAs). CatMPAs could be simply obtained by centrifugation or magnetically purified similar to MPAs, and the superior stabilities of the CatMPAs were also demonstrated. The here-presented CatMPA strategy can serve as a modular platform for enzyme immobilization, as it allows the immobilization of new targets with minimal to no construct optimization (i.e. only the addition of a SpyCatcher tag to an immobilization target would be necessary), and therefore can be advantageous when compared to existing *in vivo* immobilization methods. As evidenced by these findings, *in vivo* produced MPAs are a promising new immobilization material solely produced by biological means, yielding catalytically-active magnetic protein aggregates (CatMPAs) in a modular fashion. With our study, we thus extended the use of ferritin in biotechnology and further diversify the toolbox of (*in vivo*) enzyme immobilization methods.

## Data availability statement

All data presented in this manuscript has been included in the main text of this article and in the Supporting Information. Further inquiries can be directed to the corresponding author.

## Author Contributions

U.K. conceived and designed the study. G.O. generated all constructs, prepared MPAs and CatMPAs, performed magnetic purification and imaging over permanent magnets, activity/fluorescence assays and all other experiments unless mentioned otherwise. B.W., supervised by D.K., performed the microscopic analyses. G.O. and U.K. prepared the manuscript. All authors have read and approved the final version.

## Acknowledgments

The authors would like to thank Kira Küsters (IBG-1 Forschungszentrum Jülich) for cultivating the citrine-ferritin and citrine control constructs in her BioLector setup prior to microscopic imaging, Gabriela María Fuentes Reyes (HHU, Düsseldorf) for her assistance in cloning the soluble control constructs, and Esther Knieps-Grünhagen (IMET HHU, Düsseldorf, Forschungszentrum Jülich) for her skillful technical assistance.

