## Supporting Information for "Magnetic Protein Aggregates Generated by Supramolecular Assembly of Ferritin Cages - A Modular Strategy for the Immobilization of Enzymes"

Author affiliations


| Content | Description | Page no |
|---|---|---|
| | **Supporting Methods** | |
| | Cloning | 2 |
| | BioLector Cultivations | 2-3 |
| | Determination of MPA yields | 3 |
| | **Supporting Results** | |
| Table S1 | List of constructs | 4 |
| Table S2 | Oligonucleotides used in the study | 4 |
| Figure S1 | Self-sedimentation of crude cell extracts of MPAs | 4 |
| Table S3 | Extinction coefficients and molecular weights constructs | 5 |
| Table S4 | Yields and protein contents of the MPA constructs | 5 |
| Figure S2 | BioLector experiments with varying iron-citrate complex concentrations | 5-6 |
| Figure S3 | Crude cell extracts of constructs overproducing soluble ferritins | 6 |
| Figure S4 | Crude cell extracts of constructs overproducing MPAs | 7 |
| Table S5 | Prescreening of different CCE mixture ratios and incubation times for CatMPA formation | 7-8 |
| Table S6 | Prescreening of all bait and prey constructs tested for the CatMPA approach | 8 |
| Figure S5 | SDS-PAGE analyses | 8-10 |
| | **Supporting References** | 11 |

### Supporting Methods

#### Cloning

Control constructs containing only HuftnH/EcftnA-WT/EcftnA H34L/T64I were generated via PCR, using oligonucleotides listed in Table S2 and Citrine-HuftnH/EcftnA-WT/EcftnA H34L/T64I constructs as template for the amplification of the respective ferritin encoding genes. A 5'-NdeI site and a 3'-XhoI site were included in the oligonucleotide primers to amplify the respective ferritins with 5'-NdeI and 3'-XhoI sites, and the resulting PCR products were digested using NdeI and XhoI restriction enzymes, and ligated into similarly digested pET28a to generate the soluble HuftnH/EcftnA-WT/EcftnA H34L/T64I control constructs. The SpyTag-RADH construct was generated by using a synthetic gene containing the entire sequence of the construct and flanked by flanked by 5'-NdeI and 3'-XhoI sites, where the synthetic gene was hydrolyzed using NdeI and XhoI, and ligated to pET28a digested with the same enzymes. SpyTag-GFIL8-RADH and GFIL8-RADH-SpyTag constructs were generated via a modular construction strategy, using GFIL8-RADH construct generated earlier[1], and a synthetic gene that contained the *radh* gene only partially. As such, the synthetic gene with SpyTag-GFIL8-RADH sequence harbored a 5'-NdeI site, and contained the *radh* gene until the natural PstI site of the native *radh*. The synthetic gene was digested using NdeI and PstI, and ligated to the GFIL8-RADH vector to obtain the complete SpyTag-GFIL8-RADH construct. Similarly, the synthetic gene encoding a partial *radh* sequence (after the natural PstI site), followed by SpyTag and a 3'-XhoI site, was digested using PstI and XhoI, and ligated to GFIL8-RADH, which was identically hydrolyzed to yield the complete GFIL8-RADH-SpyTag. SpyCatcher-EcftnA H34L/T64I and SpyCatcher-Citrine-EcftnA H34L/T64I constructs were generated via the In-Fusion cloning kit (Clontech Laboratories, Inc., Takara Bio, Saint-Germain-en-Laye, France). Briefly, the synthetic gene containing the SpyCatcher sequence was used as a template for PCR using primers (Table S2) designed for In-Fusion cloning according to kit instructions, and ligation free cloning was performed using the PCR products and Citrine-EcftnA H34L/T64I vector either nicked using NdeI (to generate SpyCatcher-EcftnA H34L/T64I), or digested using NdeI and HindIII (to generate SpyCatcher-Citrine-EcftnA H34L/T64I). All SpyTag/SpyCatcher containing constructs included a linker (L) separating the genes encoding these tags from the genes encoding target proteins. All constructs generated in this study was verified by sequencing.

#### BioLector Cultivations

For Biolector (m2p-labs GmbH, Baesweiler, Germany) experiments, *E. coli* BL21(DE3) cells overproducing citrine-ferritins were initially cultivated in LB medium in shake flasks (37 °C, 130 rpm, overnight). LB precultures were used to inoculate 100 ml AI main cultures at a starting

OD600 of 0.05, and were incubated at 37 °C for 3 hours, shaking at 130 rpm. After this initial growth phase, 900 µl of the AI cultures were transferred onto 100 µl AI medium with varying iron-citrate concentrations (0 mM - 100 mM) in 48-well FlowerPlates (m2p-labs GmbH, Baesweiler, Germany), giving rise to final iron-citrate concentrations ranging between 0 mM - 10 mM in the FlowerPlates. The FlowerPlates were then immediately covered with oxygen-permeable films and transferred to the Biolector, where the expression continued for 69 hours at 15 °C, shaking at 1200 rpm. The Biolector setup was used to monitor biomass as estimated by scattered light (λex 620 nm, λem 620) and citrine fluorescence using filter sets for eYFP (λex 508 nm, λem 532 nm) during the expression for the initial assessment of varying iron-citrate complex concentrations on growth and the fluorescence of live cells.

**Determination of yields for MPA costructs**

To determine the yields of MPA constructs, the crude cell extracts of freshly lysed cells were centrifuged (15000*g*, 30 mins, 4 °C) and the supernatant was discarded. The pellet was then resuspended in the same volume of MilliQ water as the removed supernatant and centrifuged a second time (15000*g*, 30 mins, 4 °C). The supernatant of the wash was discarded, and the washed pellet was again resuspended in the same volume of MilliQ water, and the washed pellet suspension was frozen at -80 °C. The frozen pellet was weighed and then lyophilized (Christ ALPHA 1-3 LD Plus, Martin Christ Gefriertrocknungsanlagen GmbH, Osterode, Germany), ground into powder using mortar and pestles, and the lyophilized pellet was carefully weighed. The lyophilized samples were stored under an argon atmosphere at -20 °C. Protein content of the lyophilizates were determined[1] by dissolving the samples in 6 M guanidine-hydrochloride at 30 °C for 30 minutes, followed by centrifugation (7697*g*, 20 mins, room temperature), and the transfer of the supernatants to disposable cuvettes to measure absorbance at 280 nm (Cary 60 UV-Vis Spectrophotometer, Agilent, Santa Clara, USA). The protein contents of the lyophilized pellets were calculated using the theoretical extinction coefficients and molecular weights estimated from the amino acid sequences of the constructs using ProtParam tool[2] (see Table S3). Yields were calculated by dividing the amount of lyophilizate or protein obtained at the end of the process (in mg lyophilizate or mg protein) by the amount of wet cells that were lysed to obtain these lyophilizates (in g wet cells), see Table S4.

### Supporting Results

**Table S1.** List of constructs generated for the study.

| **Construct name** |
|---|
| Citrine-HuftnH[3] |
| Citrine-EcftnA-WT |
| Citrine-EcftnA H34L/T64I |
| Citrine |
| HuftnH |
| EcftnA-WT |
| EcftnA H34L/T64I[4] |
| SpyTag-GFIL8-RADH |
| GFIL8-RADH-SpyTag |
| SpyCatcher-EcftnA H34L/T64I |
| SpyCatcher-Citrine-EcftnA H34L/T64I |
| SpyTag-Citrine-EcftnA H34L/T64I |
| SpyCatcher-RADH |
| SpyTag-RADH |

**Table S2.** Oligonucleotides used in the study. The restriction sites are underlined.

| **Name** | **Sequence (5' - 3')** | **Construct generated** |
|---|---|---|
| NdeI_Citrine_fw | TATATACATATGGTGAGCAAGGGCGAGGAGCTGTTC | Soluble citrine |
| Citrine_Stop_XhoI_rev | TATATACTCGAGTTACTTGTACAGCTCGTCCATGCCG | |
| NdeI_HuftnH_fw | TATATACATATGACGACCGCATCCACCTCGCAGG | Soluble HuftnH |
| HuftnH_Stop_XhoI_rev | ATATATCTCGAGTTAGCTTTCATTATCACTGTCTCC | |
| NdeI_EcftnA_fw | TATATACATATGCTGAAACCAGAAATGATTG | Soluble EcftnA-WT<br>Soluble EcftnA H34L/T64I |
| EcftnA_XhoI_rev | ATATATCTCGAGTTAGTTTTGTGTGTCGAGGGTAGAG | |
| NdeI_SpyCatcher_R_fw | TATATACATATGGGCGCGATGGTGACCACCCTGAGCG | SpyCatcher-RADH |
| L_HindIII_rev | TATATAAAGCTTACGGCCTTCAATGCTACCGCCACCGCCGCTAC | SpyCatcher-RADH |
| NdeI_SpyCatcher_fw | AAGGAGATATACATATGGGCGCGATGGTGACCACC | SpyCatcher-Citrine-EcftnA H34L/T64I<br>SpyCatcher-EcftnA H34L/T64I |
| SpyCatcher_NdeI_rev | CCCTTGCTCACCATATGGCTACCGCCACCGCCGCTACC | SpyCatcher-Citrine-EcftnA H34L/T64I |
| SpyCatcher_HindIII_rev | GTTTCAGCATAAGCTTGCTACCGCCACCGCCGCTACC | SpyCatcher-EcftnA H34L/T64I |

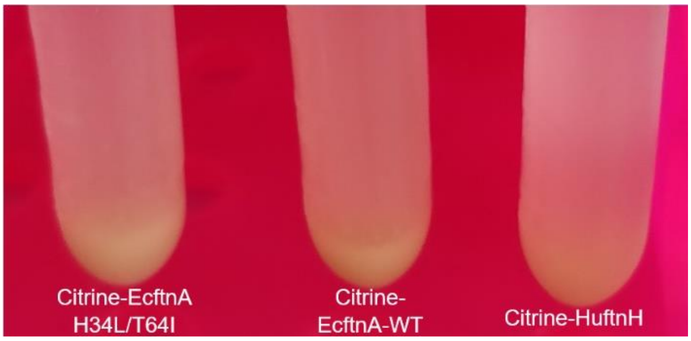

**Figure S1**. Self-sedimentation of crude cell extracts of citrine-ferritins in a test tube after 16 hours.

**Table S3.** Theoretical extinction coefficients and molecular weights of the MPA constructs derived using the amino acid sequences and Expasy ProtParam[2] (http://web.expasy.org/protparam)

| Construct name | Extinction coefficient ($M^{-1}$ $cm^{-1}$) | Molecular weight (Da) |
|---|---|---|
| Citrine-HuftnH | 42540 | 49.6 |
| Citrine-EcftnA-WT | 47915 | 47.8 |
| Citrine-EcftnA H34L/T64I | 47915 | 47.8 |

**Table S4.** Yields and protein contents of the MPA constructs generated in the study.

| Construct name | Yield (g lyophilizate / 100 g wet cells) | | Yield (mg protein / g wet cells) | | Protein content of lyophilizate (%) | |
|---|---|---|---|---|---|---|
| | Mean | SE | Mean | SE | Mean | SE |
| Citrine-HuftnH | 4.7 | 0.2 | 36.5 | 3.6 | 76.6 | 4.6 |
| Citrine-EcftnA-WT | 3.8 | 0.4 | 22.2 | 3.0 | 57.5 | 2.3 |
| Citrine-EcftnA H34L/T64I | 3.8 | 0.5 | 29.1 | 3.6 | 76.2 | 1.7 |

SE represents standard error of the mean derived from at least three biological replicates with three technical replicates each. Protein contents of lyophilizates were calculated using the theoretical extinction coefficients and molecular weights listed in Table S3.

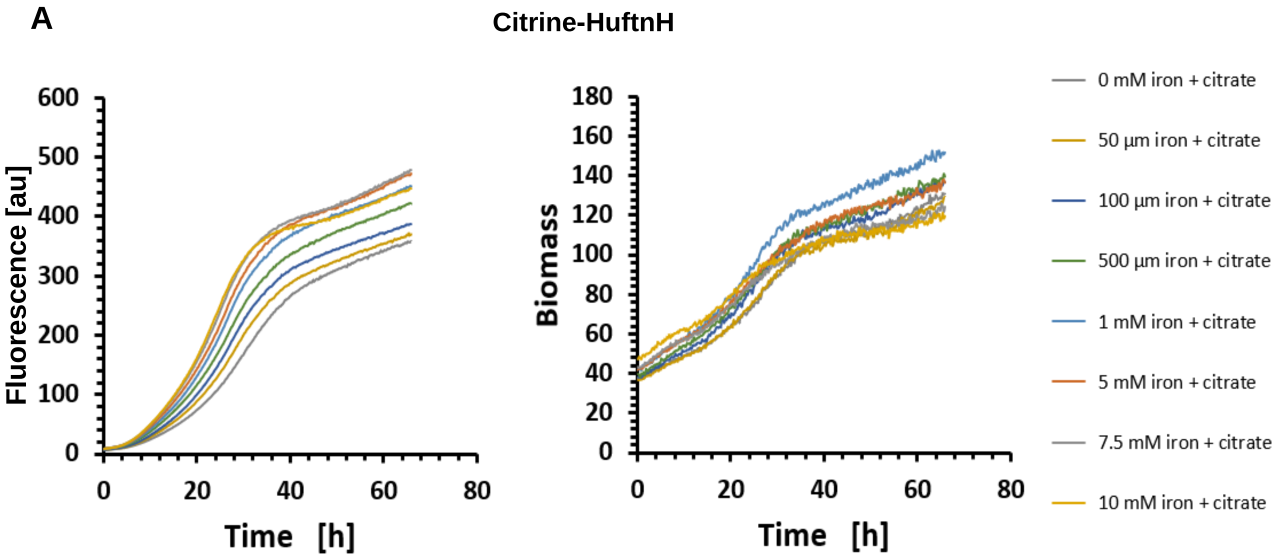


**Figure S2.** BioLector experiments depicting citrine fluorescence (left) and biomass (right) during expression with varying iron-citrate complex concentrations for **(A)** Citrine-HuftnH, **(B)** Citrine-EcftnA-WT and **(C)** Citrine-EcftnA H34L/T64I. Citrine-ferritins that were not supplemented with iron displayed the lowest fluorescence intensity for all three constructs, which was followed by 50 µM, 100 µM and 500 µM supplementations, indicating that iron concentration has a marked effect on the proper maturation of the citrine-ferritin fusion proteins. Supplementation of 5 mM or more of iron citrate complex had a negative impact on growth for Citrine-EcftnA-WT and Citrine-EcftnA-H34L/T64I, and the same effect was observed for supplementation of 7.5 mM or more iron for Citrine-HuftnH construct. Therefore, 1 mM was chosen as a suitable concentration for iron supplementation and was used for the cultivation of all strains.

**Figure S2 (continued)**

**B** **Citrine-EcftnA-WT**

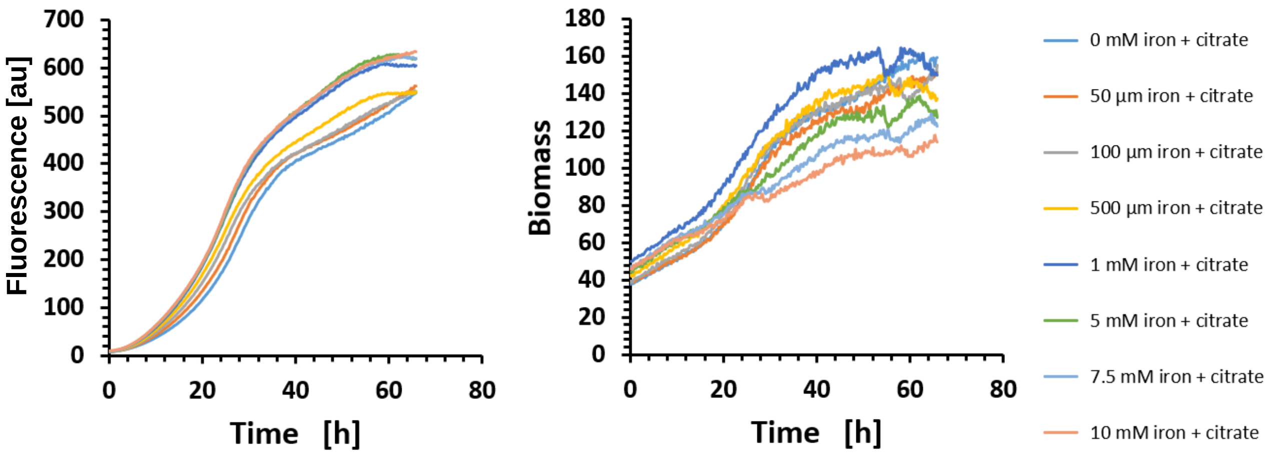


**C** **Citrine-EcftnA H34L/T64I**

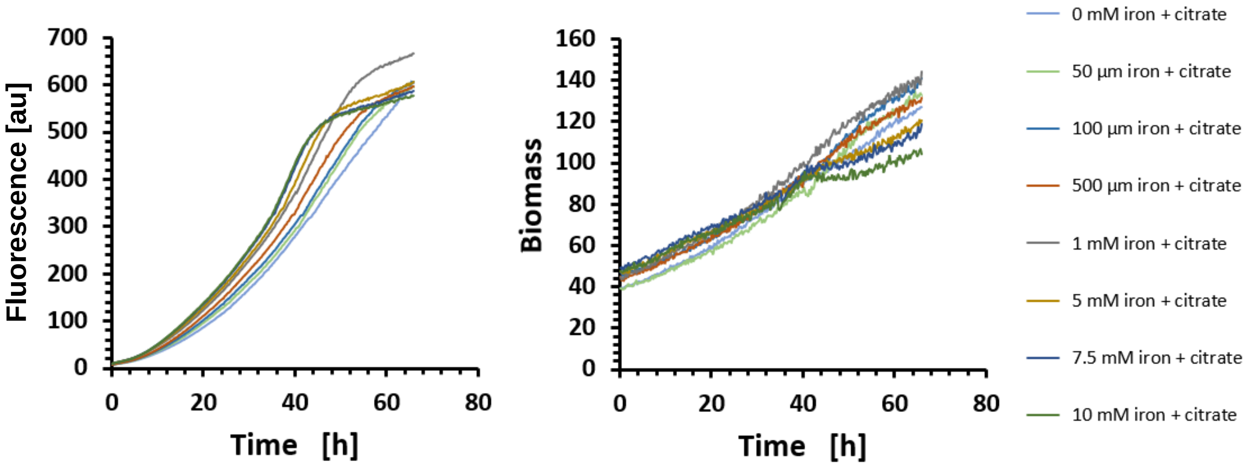


**HuftnH** **EcftnA-WT** **EcftnA H34L/T64I**

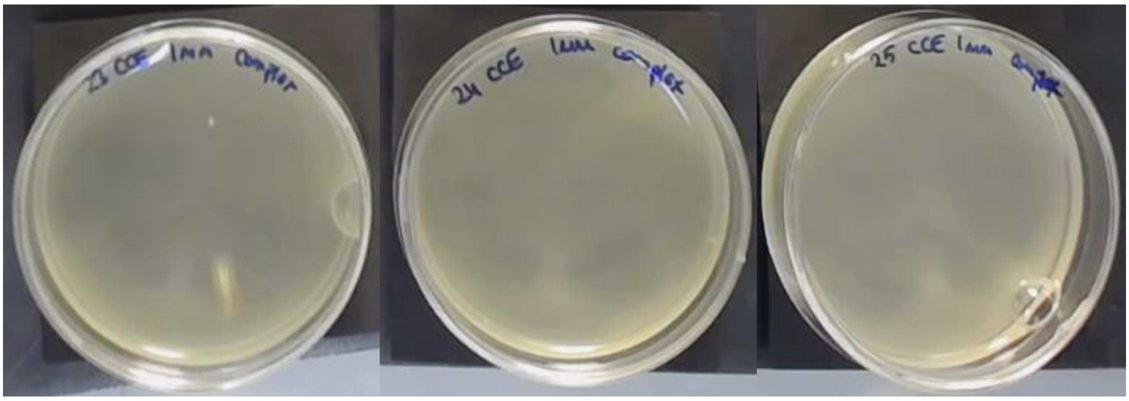

**Figure S3.** Crude cell extracts (CCEs) of constructs overproducing soluble ferritins. Mini petri dishes containing CCEs were placed over permanent neodymium ring magnets arranged in a 2x2 grid covered with a black paper. The CCEs were mixed with OptiPrep density gradient medium mixture (17% Optiprep) and imaged after 69 hours. The contrast of all three images were increased by 20%.

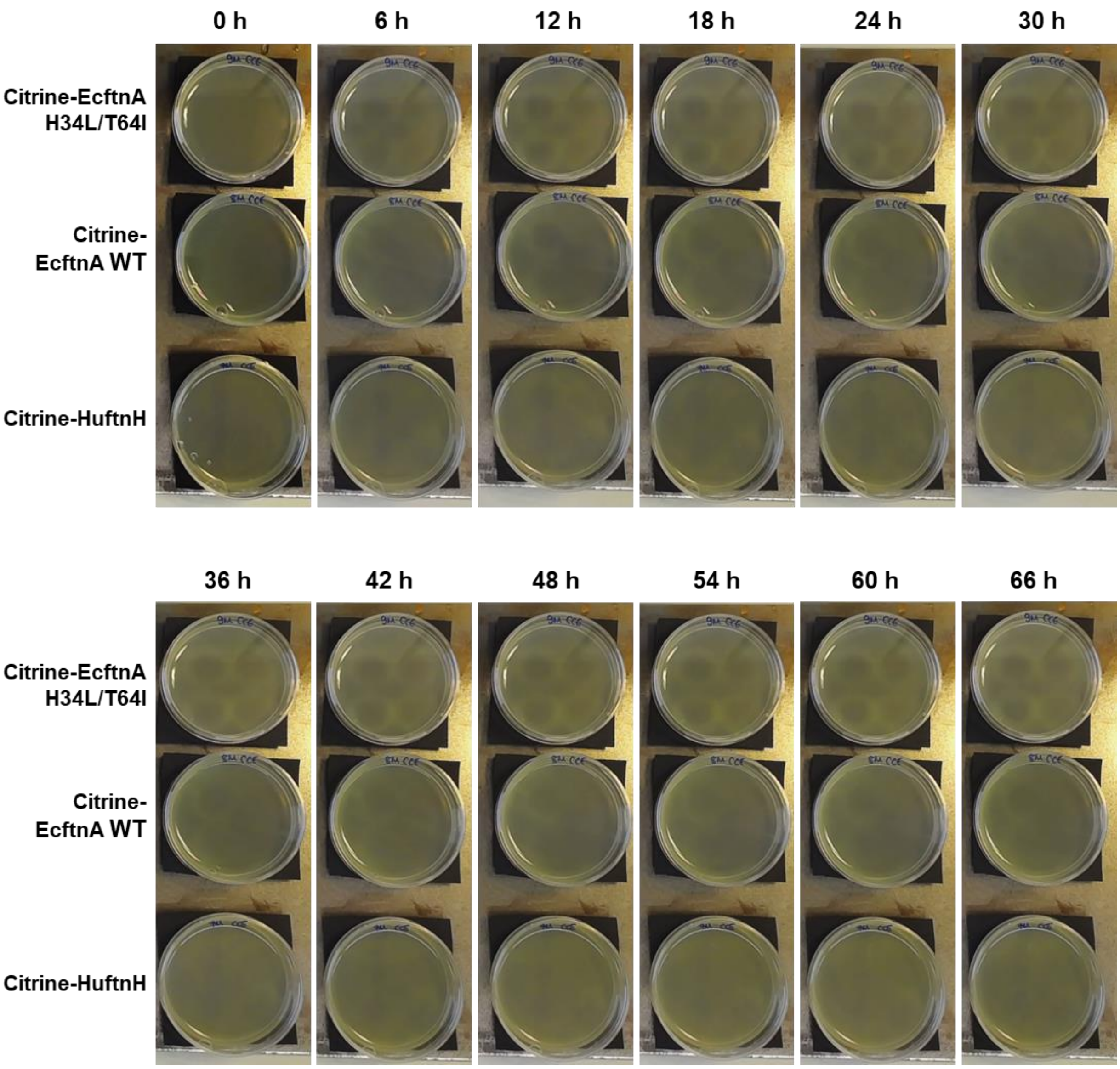


**Figure S4.** Crude cell extracts (CCEs) of Citrine-HuftnH, Citrine-EcftnA WT and Citrine-EcftnA H34L/T64I MPA constructs placed over permanent neodymium ring magnets. The CCEs were mixed with OptiPrep density gradient medium mixture (17% Optiprep) and visualized up to 69 hours, where images show the pattern progression in 6 hour intervals. The contrast of all three images were increased by 20%.

**Table S5.** Prescreening of SpyTag-Citrine-EcftnA H34L/T64I (bait) and SpyCatcher-RADH (prey) constructs for the initial assessment of the effect of different mixing ratios and incubation times on immobilization success, judged by the RADH activity distribution of the cell fractions.

| **Mixture ratio Bait:Prey** (v/v) | **Incubation time (minutes)** | **Relative RADH Activity (%)** | | |
|---|---|---|---|---|
| | | **CCE** | **S** | **P** |
| 1:1 | 30 | 100 | 65.4 | 35.1 |
| 1:1 | 60 | 100 | 75.3 | 38.2 |
| 1:1 | 90 | 100 | 67.9 | 35.6 |
| 1:10 | 30 | 100 | 73.0 | 34.1 |
| 1:20 | 30 | 100 | 77.7 | 25.4 |
| 5:1 | 30 | 100 | 74.1 | 33.3 |
| 10:1 | 30 | 100 | 76.3 | 27.6 |

The crude cell extracts (CCEs) of bait and prey constructs were either mixed in a 1:1 ratio and incubated for different durations, or were incubated for 30 minutes but mixed in a different ratio. For all cases, after bait and prey constructs were cultivated and lysed separately, their CCEs were mixed and incubated at 25 °C, fractionated to yield soluble (S) and insoluble fractions, and the insoluble fractions were washed to obtained washed pellets (P). Each fraction was compared to the total RADH activity of the mixed CCE for each case (set to 100%).

**Table S6**. Prescreening of all bait and prey constructs used for the CatMPA approach for the initial assessment of RADH activity distribution and purification efficiencies.

| **Names of constructs mixed (1:1 v/v)** | | **Relative RADH Activity of bait + prey mixture (%)** | | | **Relative RADH Activity of prey (%)** | | | **Purification efficiency (%)** |
|---|---|---|---|---|---|---|---|---|
| **Bait** | **Prey** | **CCE** | **S** | **P** | **CCE** | **S** | **P** | **Bait + Prey** |
| SpyTag-Citrine-EcftnA H34L/T64I (*) | SpyCatcher-RADH | 100 | 65.4 | 35.1 | 100 | 89.5 | 12.8 | 9.4 - 18.4** |
| SpyCatcher-Citrine-EcftnA H34L/T64I (*) | SpyTag-RADH | 100 | 91.5 | 8.3 | 100 | 97.2 | 2.6 | n.a. |
| SpyCatcher-EcftnA H34L/T64I | SpyTag-GFIL8-RADH (*) | 100 | 75.6 | 30.9 | 100 | 58.8 | 40.2 | 5.1 |
| SpyCatcher-EcftnA H34L/T64I | GFIL8-RADH-SpyTag (*) | 100 | 73.9 | 24.2 | 100 | 72.7 | 27.7 | 97.4 |
| Citrine-EcftnA H34L/T64I (*) | SpyCatcher-RADH | 100 | 99 | 0.89 | 100 | 97.8 | 1.66 | n.d. |
| SpyTag-Citrine-EcftnA H34L/T64I | Soluble RADH | 100 | 97.1 | 1.43 | 100 | 98.3 | 0.92 | n.d. |

Crude cell extracts (CCE) were fractionated to yield soluble (S) and insoluble fractions, and the insoluble fractions were washed to obtained washed pellets (P). For bait + prey, bait and prey constructs were cultivated and lysed separately, their CCEs were mixed in a 1:1 ratio (v/v) and incubated at 25 °C for 30 minutes, fractionated and washed to yield S and P fractions of the CCE mixture in the same manner. All fractions were compared to the total RADH activity of the CCE they were fractionated from (mixed CCE for bait+prey) for each case (set to 100%). For each pair, the insoluble partner is marked with an asterisk (*). Purification efficiencies were calculated using mixed CCE fractions, based on relative citrine fluorescence or relative RADH activity (**) of the magnetic fraction (MG), compared to that of the washed pellet (P2, set to 100%), according to the formula; $\boldsymbol{Purification\ Efficiency}$ **[%]** = ($Fluorescence\ of\ MG$ [$AU$]) / ($Fluorescence\ of\ P$ [$AU$]) $x$ 100. For the bait-prey mixtures with a single purification efficiency value, the calculation is based on RADH activity. The last two rows show the negative control pairs (Citrine-EcftnA H34L/T64I + SpyCatcher-RADH, and SpyTag-Citrine- EcftnA H34L/T64I + Soluble RADH) assayed to exclude the effect of hydrophobic interactions for CatMPA formation.

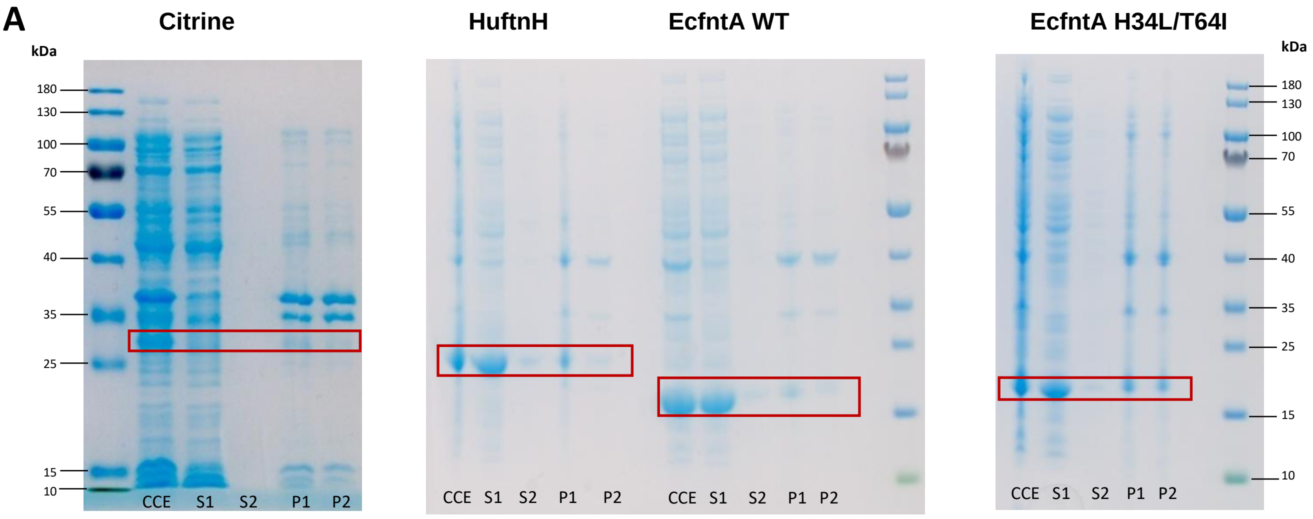


**Figure S5**. SDS-PAGE analyses of cell fractions from all MPA, CatMPA, and control constructs. In each panel, the expected size of the fusion protein is indicated via the red rectangles. CCE: crude cell extract. S1: supernatant. S2: supernatant of the wash step. P1: unwashed pellet. P2: washed pellet (MPA or CatMPA fraction). For CatMPAs, the magnetic purification fractions are denoted as follows; NM: nonmagnetic fraction, W1: first wash, W2: second wash, MG: magnetic fraction. **Panel A** depicts the MPA control constructs containing either only citrine, or only ferritin. From left to right: Citrine (27 kDa), HuftnH (21.2 kDa), EcftnA WT (19.4 kDa), EcfntA H34L/T64I (19.4 kDa). **Panel B** depicts the cell fractions of MPA constructs. From left to right: Citrine-HuftnH (49.6 kDa), Citrine-EcfntA WT (47.8 kDa kDa), Citrine-EcfntA H34L/T64I (47.8 kDa). Panels C, D, E and F depict CatMPA constructs, where EcftnA refers to the mutant ferritin for each case (H34L/T64I). In each panel, the cell fractions of lone SpyTag or SpyCatcher containing bait or prey constructs are shown, together with the fractions of the mixed crude cell extracts of each relevant pair. The mixed crude cell extracts were obtained by mixing the CCEs of the corresponding bait/prey constructs in a 1:1 ratio followed by a 30 minute incubation to allow the SpyTag/SpyCatcher reaction to take place, as described in text (see Methods). Afterwards where the CCE mixtures were fractionated, and further purified using magnetic columns**. Panel C:** From left to right: SpyTag-Citrine-EcftnA (bait, 50.2 kDa), SpyCatcher-RADH (prey, 40.6 kDa), SpyTag-Citrine-EcftnA + SpyCatcher-RADH (bait+prey, 90.8 kDa). **Panel D:** From left to right: SpyCatcher-EcftnA (bait,32.8 kDa), SpyCatcher-Citrine-EcftnA (bait, 61.1 kDa) SpyTag-GFIL8-RADH (prey, 31.9 kDa), GFIL8-RADH-SpyTag (prey, 31.9 kDa). **Panel E:** From left to right: SpyTag-GFIL8-RADH + SpyCatcher-EcftnA (bait+prey, 64.7 kDa), GFIL8-RADH-SpyTag + SpyCatcher-EcftnA (bait+prey, 64.7 kDa). **Panel F:** From left to right: SpyCatcher-Citrine-EcftnA (bait, 61.1 kDa), SpyTag-RADH (prey, 29.4 kDa). For the pair depicted in panel F (SpyCatcher-Citrine-EcftnA and SpyTag-RADH), the bait construct almost completely lost its ability to form insoluble MPAs by the addition of the larger SpyCatcher tag at the N-terminus, as evidenced by the relative fluorescence assay (91.5% soluble, Table S6).Upon mixing of the CCEs, the pellet displayed low RADH activity (8.3%; Table S6). Therefore this bait/prey combination was not assayed further via SDS-PAGE analyses.

**Figure S5 (continued)**

**B** **Citrine-HuftnH** **Citrine-EcfntA WT** **Citrine-EcfntA H34L/T64I**

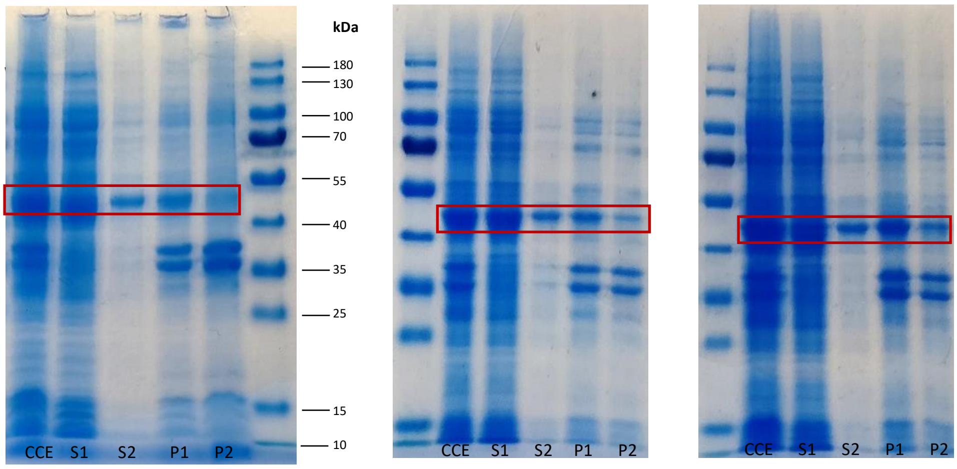


**C** **SpyTag-Citrine-EcftnA - SpyCatcher-RADH** **SpyTag-Citrine-EcftnA + SpyCatcher-RADH**

**(Bait)** **(Prey)** **(Bait + Prey)**

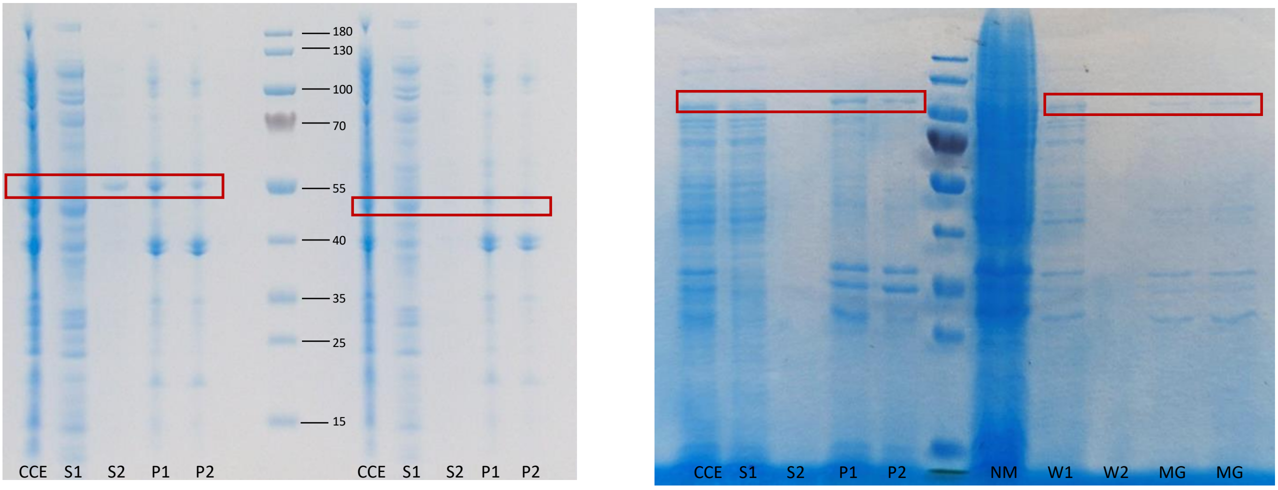

**Figure S5 (continued)**

**D**

**SpyCatcher-EcftnA (Bait)** **SpyCatcher-Citrine-EcftnA (Bait)**

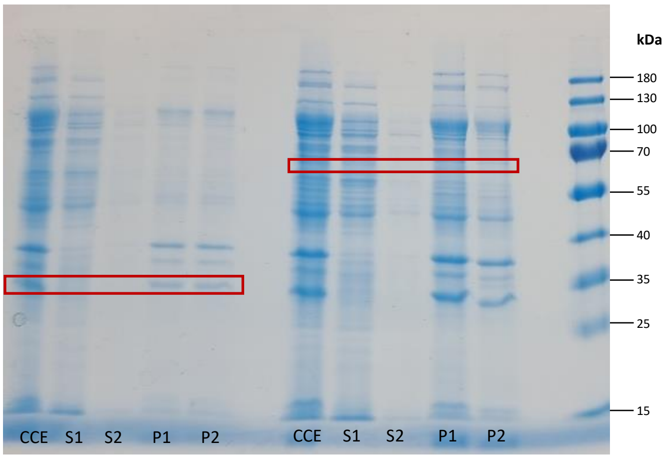


**SpyTag-GFIL8-RADH (Prey)** **GFIL8-RADH-SpyTag (Prey)**

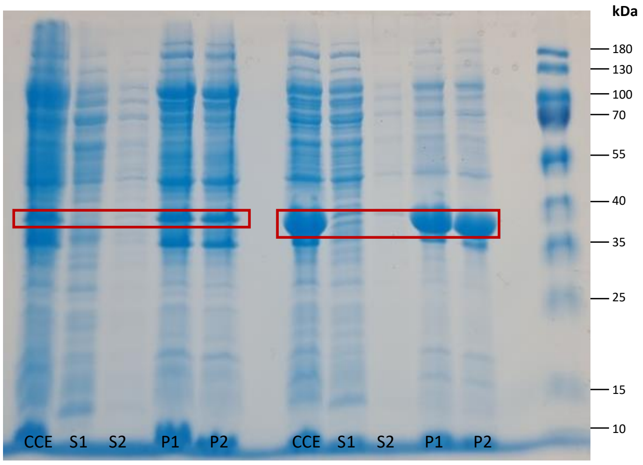


**E**

**SpyTag- GFIL8-RADH + SpyCatcher-EcftnA (Bait+Prey)**

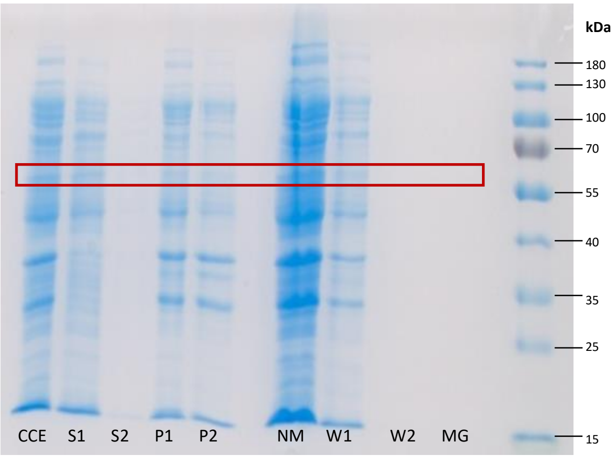


**GFIL8-RADH-SpyTag + SpyCatcher-EcftnA (Bait+Prey)**

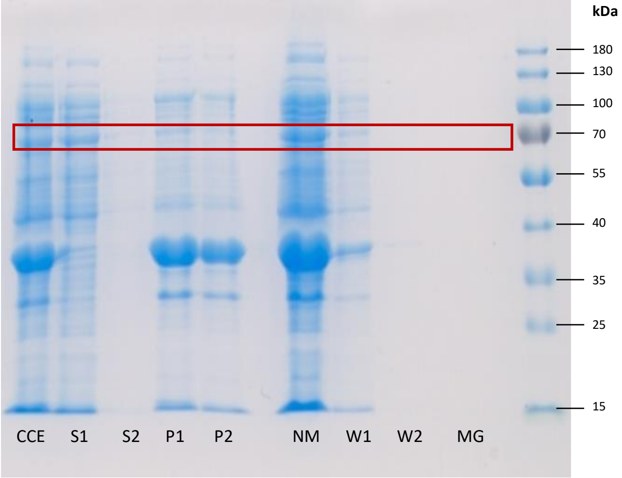


**F**

**SpyCatcher-Citrine- EcftnA (Bait)**

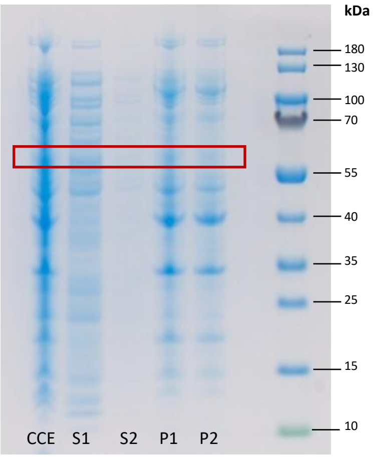


**SpyTag-RADH (Prey)**

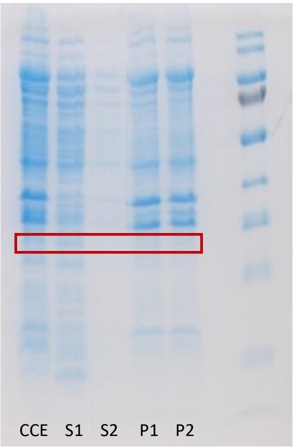
